# IMC10 and LMF1 mediate membrane contact between the mitochondrion and the inner membrane complex in *Toxoplasma gondii*

**DOI:** 10.1101/2022.04.01.486766

**Authors:** Rodolpho Ornitz Oliveira Souza, Kylie N. Jacobs, Gustavo Arrizabalaga

## Abstract

The single mitochondrion of *Toxoplasma gondii* is highly dynamic, being predominantly in a peripherally distributed lasso-shape in intracellular parasites and collapsed in extracellular ones. The peripheral positioning of the mitochondrion is associated with apparent contacts between the mitochondrion membrane and the parasite pellicle. The outer mitochondrial membrane-associated protein LMF1 is critical for the correct positioning of the mitochondrion, and in its absence, intracellular parasites fail to form the lasso-shaped mitochondrion. To identify other proteins that participate in tethering the parasite’s mitochondrion to the pellicle, we performed a yeast two-hybrid screen for LMF1 interactors. We identified 70 putative interactors, six of which are known to localize to the apical end of the parasite, two to the mitochondrial membrane, and three localize to the inner membrane complex (IMC), a component of the parasite pellicle. Using reciprocal immunoprecipitation and proximity ligation assays, we confirmed the interaction of LMF1 with the pellicle protein IMC10, with a hypothetical protein known to be part of the conoid, and with an ATPase-Guanylyl Cyclase. Conditional knockdown of IMC10 does not affect parasite viability but severely affects mitochondrial morphology in intracellular parasites and mitochondrial distribution to the daughter cells during division. In effect, IMC10 knockdown phenocopies disruption of LMF1, suggesting that these two proteins define a novel membrane tether between *Toxoplasma*’s mitochondrion and the inner membrane complex.

IMPORTANCE

*Toxoplasma gondii* is an opportunistic parasite that can cause life-threatening disease in immunocompromised patients and those infected congenitally. As current therapies against this parasite can be poorly tolerated and are not effective against the latent stage of the parasite, there is an urgent need to identify new drug targets. The single mitochondrion of this parasite is a validated drug target, but little is known about the machinery that controls its division and structure, information that would be critical for a thorough exploration of the mitochondrion as a drug target. We have identified parasite-specific proteins that are essential to maintain the normal structure of the mitochondrion. We have discovered a complex of two proteins that tether the mitochondrion to the periphery of the parasite. Loss of this connection results in changes in mitochondrial morphology and cell division defects. Our results provide important insight into the molecular mechanisms regulating *Toxoplasma* mitochondrial morphology.

## INTRODUCTION

*Toxoplasma gondii* is a highly successful intracellular pathogen that belongs to the phylum Apicomplexa (Black and Boothroyd, 2000) and is the causative agent of toxoplasmosis (Hill et al., 2005). This parasite can infect any nucleated cell in a plethora of warm-blooded animals. It is estimated that approximately 30% of the human population might be infected with *Toxoplasma* (Pappas et al., 2009). Although most infections are asymptomatic, toxoplasmosis is a severe problem for immunosuppressed patients (Porter and Sande, 1992) and in congenital infections (Khan and Khan, 2018). Drugs against these pathogens are limited, often toxic, and, resistance is a serious challenge. Thus, the discovery of novel therapeutics is a priority.

A unique feature of this parasite is the presence of a single tubular mitochondrion, which is essential for parasite survival and a validated drug target. *Toxoplasma*’s mitochondrion is highly dynamic, showing different morphologies during the parasite’s propagation cycle and in response to stress factors (Charvat and Arrizabalaga, 2016; Jacobs et al., 2020; Ovciarikova et al., 2017). When the parasite is within a host cell, the mitochondrion is in a lasso shape, distributed along the periphery of the cell and adjacent to the parasite’s pellicle. When in the extracellular environment, the mitochondrion collapses towards the apical end of the parasite. During this transition, some of the parasites can present an intermediate stage morphology called “sperm-like” (Ovciarikova et al., 2017). As soon as the parasite re-enters a cell, the mitochondrion recovers the lasso shape. It has been observed that when in the lasso shape, the mitochondrion has patches of its membrane in close proximity to the parasite’s pellicle, reminiscent of membrane contact sites (MCS) (Ovciarikova et al., 2017).

*Toxoplasma*’s pellicle is composed of the parasite plasma membrane, the inner membrane complex (IMC), which consists of a series of flattened membrane sacs, and a supporting network of intermediate filaments. Contact between the mitochondrion and the elements of the pellicle is also observed during cell division. *Toxoplasma* divides by a specialized process called endodyogeny, where two daughter cells emerge within the mother cell (Hu et al., 2002). During this process, the IMC serves as a scaffold for the segregation and division of parasite organelles (Nishi et al., 2008). As there is only one mitochondrion per parasite, its division is tightly coordinated with the division of the rest of the parasite (Nishi et al., 2008; Verhoef et al., 2021). As the two nascent IMCs form during endodyogeny, the mitochondrion develops extensions along its length, which continue to grow as the daughter IMCs elongate. The branching mitochondrion is excluded from the daughter parasites until the latest stage of division, at which point mitochondrial branches enter the developing daughters moving along the IMC scaffold as the nascent parasites emerge from the mother cell. (Nishi et al., 2008; Ovciarikova et al., 2017). Thus, the mitochondrion is highly dynamic as the parasite moves in and out of cells and during parasite division. While the dynamics of the mitochondrion have been well described, our understanding of the mechanisms and the proteins that drive them remains vague.

Apicomplexan organisms must have evolved new ways to divide and distribute the mitochondrion, given the fact that most of the proteins involved in mitochondrial fission and fusion found in opisthokonta are not present in the genome of these organisms (Verhoef et al., 2021; Voleman and Dolezal, 2019). Recently our laboratory reported that a homolog of the yeast Fission Protein 1 (Fis1) is located at the outer mitochondrial membrane (OMM), but it is not essential for mitochondrial division or parasite survival *in vitro* (Jacobs et al., 2020). Interestingly, a dominant-negative version of this protein affects mitochondrial shape and positioning. While investigating the proteins that interact with Fis1, we found a coccidian-specific protein, TGGT1_265180, that localizes to the outer mitochondrial membrane (OMM). We have named this protein the Lasso Maintenance Factor 1 (LMF1) due to the remarkable phenotype observed in its absence. Parasites lacking LMF1 are not able to form a lasso-shaped mitochondrion. Instead, the organelle is either collapsed or sperm-like in intracellular parasites, suggesting that this protein is critical for mitochondrial shaping and positioning. In addition to the mitochondrial morphology phenotype, lack of LMF1 affects parasite fitness and mitochondrial segregation into daughter cells during division, which results in amitochondriate parasites and extracellular mitochondrial material (Jacobs et al., 2020).

LMF1 has no lipid binding or transmembrane domain, which opens questions about how this protein is regulating mitochondrial shape. We previously determined that protein-protein interaction between LMF1 and Fis1 is required for the association of LMF1 with the mitochondrion and its function in maintaining the normal morphology of the mitochondrion. We hypothesize that LMF1 interacts with other proteins that facilitate the contact between the OMM and the parasite pellicle. In this work, we show that, indeed, LMF1 interacts with proteins located in the pellicle and the apical complex. Among these interactors, we found that the inner membrane protein IMC10 interacts with LMF1 to regulate mitochondrial shape and positioning. Inducible knockdown of IMC10 leads to loss of lasso shape and other mitochondrial abnormalities that phenocopy the effects of LMF1 deletion.

## RESULTS

### Phylogenetic analysis of LMF1

InterPro predictions for structured domains in the LMF1 sequence show that this protein might be organized in three different domains (N-terminal, middle and C-terminal domains) (Figure 1A). InterPro predicts two intrinsically disordered domains (aa104-181 and aa319-376). The presence of these putative disordered domains reinforces the hypothesis that LMF1 interacts with other proteins. Using a reciprocal BLAST querying of genomic sequences with the *Toxoplasma* LMF1 sequence, it was possible to recover 37 orthologues, distributed into two phyletic groups: alveolates and cryptophyte. We identified homologs present in other coccidia (e.g., *Neospora*, *Sarcocystis,* and *Hammondia*) and eimeriids, but not in *Cryptosporidium* and haemosporidians such as *Plasmodium* (Figure 1B). Our phylogenetic analysis shows that LMF1 appeared very early in the cryptist heterotrophic algae *Guillardia theta* (26% identity with *Toxoplasma* LMF1). This organism possesses a very divergent sequence that shares homology with LMF1. The chromerid *Chromera velia*, a phototrophic organism related to the apicomplexans, also encodes an LMF1 homolog that shares 28% identity in amino acid composition with the *Toxoplasma* protein (Figure 1C).

**Figure 1.**
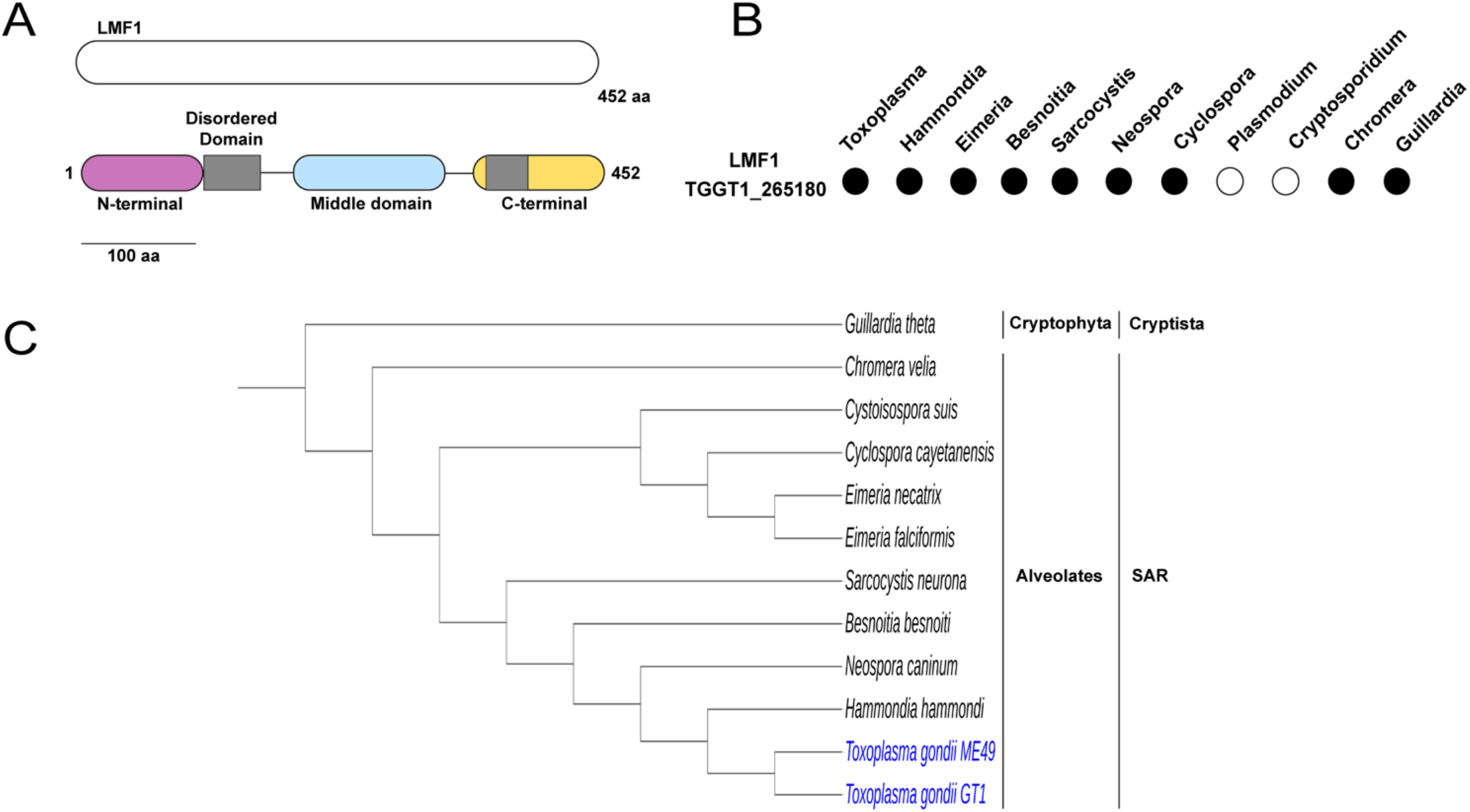
Domains and phylogenetic analysis of LMF1. A) Predicted domain architecture of LMF1. Schematic of LMF1 highlighting the three domains based on InterPro and MobiDB. Magenta: N-terminal, cyan: middle domain, yellow: C-terminal domain, and grey: predicted disordered domains. B) List of apicomplexan parasites with LMF1 orthologs. Black filled circle means the presence of an orthologue, and a white-filled circle means the absence of an LMF1 orthologue. C) Cladogram of LMF1 and LMF1 related sequences from other organisms. See materials and methods for accession numbers.

### LMF1 interacts with proteins localized to different cell compartments

To identify potential interactors of LMF1, we employed a yeast two-hybrid (Y2H) interaction screen. For this assay, we used full-length LMF1 as bait and analyzed over 95 million interactions using a *Toxoplasma* cDNA library. This screen yielded 257 positive clones, from which 69 putative interactors were identified (Dataset 1). These putative interactors were categorized based on the likelihood of interaction with LMF1 using the Predicted Biological Score (PrBS), which ranks interactors from A (highest confidence score) to D (lowest confidence score) (Dataset 1) (Formstecher et al., 2005; Fromont-Racine et al., 1997). In total, there were 4 A interactors, 6 B, 8 C, and 51 D. As to narrow this list to those that are most likely to interact with LMF1, we considered their cellular localization. Given that LMF1 is localized to the outer mitochondrial membrane and that the mitochondrion has contacts with the pellicle in intracellular parasites and the apical end in extracellular parasites, we narrowed our list to those known to be localized to either the mitochondrion, the pellicle, and the apical end of the parasite. The localization was based on a published spatial proteomics analysis (Barylyuk et al., 2020). This analysis resulted in a list of 15 proteins with three located at the pellicle, seven apical, and two mitochondrial (Table 1). Interestingly, based on the predicted CRISPR score, most of the putative interactors are fitness conferring during in vitro culture (Sidik et al., 2016).

**Table 1.** Candidate LMF1 interactors. Listed are proteins identified by the yeast two-hybrid (Y2H) screen that localize to either the pellicle (pink), apical end (yellow), or mitochondrion (light blue). Included are the Gene ID, the gene annotation, the global PBS score for the likelihood of interaction in Y2H, and the fitness score from the genome-wide CRISPR screen (Sidik et al., 2016).

| ToxoDB Gene ID | Product Description | PrBS | CRISPR score |
| --- | --- | --- | --- |
| **TGME49_230210** | **Alveolin domain-containing intermediate filament IMC10** | A | -4.7 |
| TGME49_235470 | Myosin A | B | -3.09 |
| TGME49_313380 | ILP1 | C | -4.7 |
| **TGME49_254370** | **Guanylyl cyclase** | B | -3.56 |
| TGME49_243250 | Myosin H | D | -3.94 |
| TGME49_244470 | Hypothetical protein | D | -4.21 |
| **TGME49_246720** | **Hypothetical protein** | D | 0.24 |
| TGME49_252880 | Hypothetical protein | D | -2.09 |
| TGME49_273560 | Kinesin heavy chain, putative | D | -0.94 |
| **TGME49_289990** | **Hypothetical protein** | D | -0.65 |
| **TGME49_213670** | **Hypothetical protein** | D | -3.88 |
| **TGME49_231930** | **Hypothetical protein** | D | -1.24 |

The putative interactor with the highest confidence score was TGGT1_230210, also known as IMC10, which has been previously identified as a component of the IMC (Anderson-White et al., 2011). Interestingly, when we immunoprecipitated (IP) LMF1-HA, we identified IMC10 by mass spectrometry among the 18 identified proteins that had at least five peptides in the experimental IP and none in the control IP (Supplemental Table S1). To further confirm the interaction with IMC10 and explore other proteins identified in the Y2H screen, we introduced a C-terminal Myc epitope tag to putative interactors in the cell line expressing the HA epitope-tagged LMF1 (LMF1-HA). For this analysis we selected six proteins from the Y2H interactors list: IMC10, TGGT1_246720, ATPase-Guanylyl Cyclase (TGGT1_254370), TGGT1_213670, TGGT1_289990 and TGGT1_231930). TGGT1_246720 was selected because it had been previously identified as an interactor of the OMM protein Fis1, which also interacts with LMF1 (Jacobs et al., 2020). The ATPase-Guanylyl Cyclase, which has been shown to be localized to the parasite’s apical complex (Koreny et al., 2021; Long et al., 2017), was selected for analysis given its potential regulatory role. TGGT1_289990 was selected because it is a predicted apical protein present only in coccidia (such as LMF1) and contains several disordered domains and a coiled-coiled domain, while TGGT1_213670 and TGGT1_231930 were selected due to their localization to the mitochondrion.

Once dual tagged lines were established, we performed reciprocal immunoprecipitation assays using both HA and Myc-conjugated magnetic beads (Fig. 2A). Using this technique, we determined that immunoprecipitation of LMF1 with HA beads brought down IMC10, TGGT1_246720, and ATPase-Guanylyl cyclase, confirming these interactions. Similarly, we detect LMF1 when we IP either IMC10 or ATPase-Guanylyl cyclase (Fig. 2A). On the other hand, the interactions with TGGT1_289990, TGGT1_213670, and TGGT1_231930 could not be confirmed through immunoprecipitation of either LMF1 or the putative interactor (Supplemental Figure S1A).

**Figure 2.**
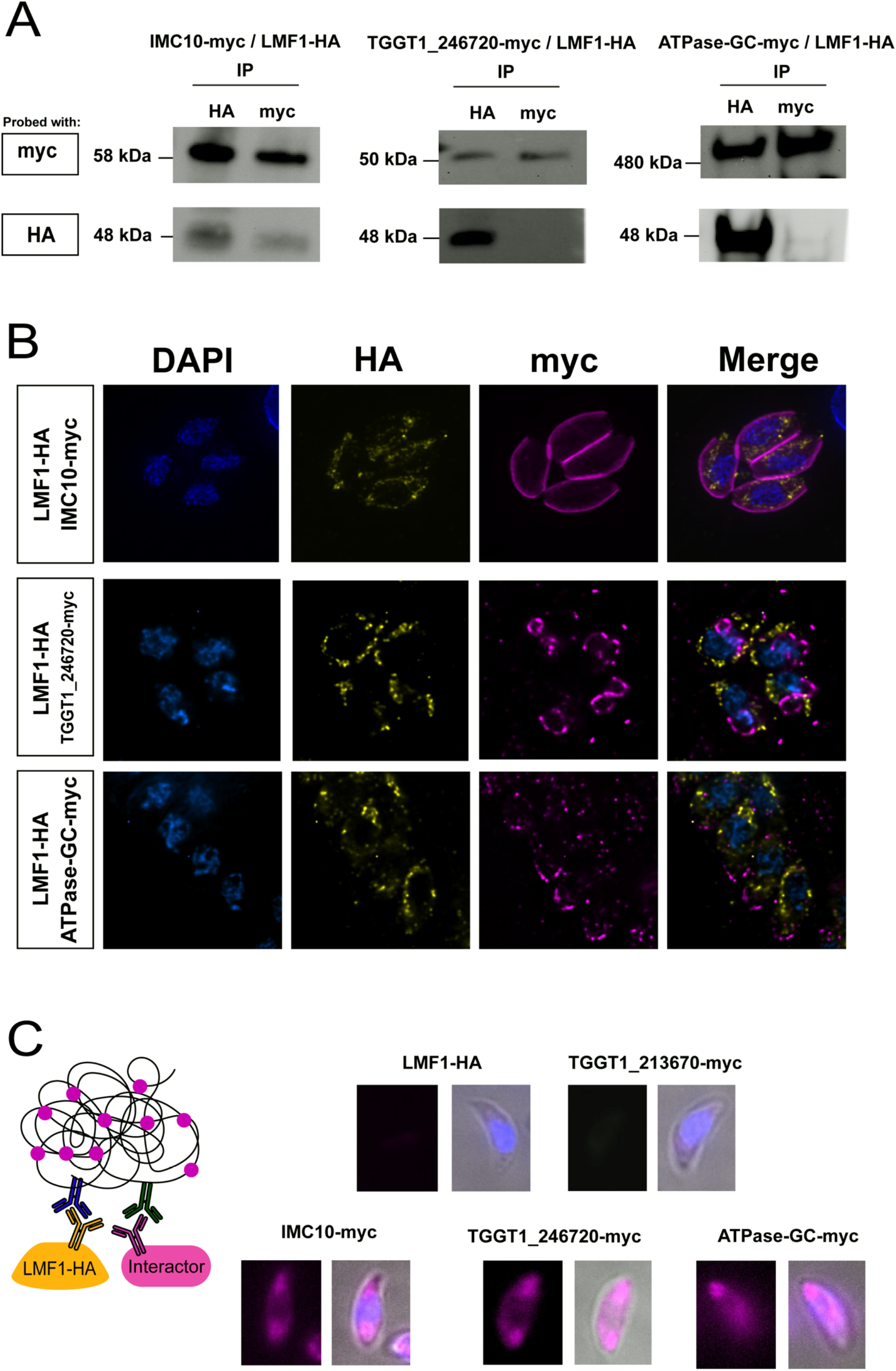
Characterization of LMF1 interactors. To investigate the localization of LMF1 interactors, we introduced sequences encoding an N-terminal Myc tag to the endogenous locus in the parasite strain expressing an HA-tagged LMF1. A) Reciprocal co-immunoprecipitation of putative LMF1 interactors was performed for the strains expressing LMF1-HA and either IMC10-Myc, TGGT1_246720-Myc, or ATPase-GC-Myc. For each of the three dually tagged parasite strains, proteins were immunoprecipitated with either anti-HA or anti-Myc conjugated beads and probed with either Myc (for the interactor) and for HA (for LMF1). B) Intracellular parasites expressing the Myc tagged versions of IMC10, TGGT1_246720 and ATPase-GC were stained for HA (yellow) and Myc (magenta). C). On the left, a schematic representation of the Proximity Ligation Assay (PLA) approach is depicted. A signal is only expected when the two proteins labeled with the primary antibodies are in proximity of each other. Images show the result of PLA for the strain expressing only LMF1-HA and the dually tagged strains. TGGT1_213670 serves as a control as it was shown to not be an interactor of LMF1 by reciprocal IP (Supplemental Figure S1).

Immunofluorescence assays (IFA) of the dual tagged lines confirmed the previously described localization of IMC10 to the IMC (Anderson-White et al., 2011) and ATPase-Guanylyl cyclase to the apical end (Brown and Sibley, 2018) (Fig. 2B). TGGT1_246720 was previously determined to localize to the conoid (Koreny et al., 2021; Long et al., 2017), which we confirmed by IFA, but we also detect this protein at the budding daughter cells, in a similar pattern as the growing IMC (Fig. 2B). As for those that did not interact with LMF1 based on IP, TGGT1_213670 and TGGT1_231930 appear to be in the mitochondrion, while TGGT1_289990 showed a punctate staining pattern throughout the parasite (Supplemental Figure S1B).

As a complementary confirmation of the interactions, we performed a proximity ligation assay (PLA) (Alam, 2018), which has been validated for use in *Toxoplasma* (Long et al., 2017; Mallo et al., 2021). We observed specific amplification of signal for all three interactors that had been confirmed by co-IP (Fig. 2C). Interestingly, the amplification of the signal was not corresponding to the shape of the mitochondrion but followed the shape of the parasite. In contrast, when we applied PLA with TGGT1_213670, a protein that was determined not to interact with LMF1 based on co-IP, no signal amplification was detected. As an additional control, we used our parental LMF1-HA cell line with both antibodies, which, as expected, did not result in amplification. Together, these results show that IMC10, TGGT1_246720, and TGGT1_254370 appear to be true LMF1 interactors within the parasite.

### Ultrastructure Expansion Microscopy (U-Ex) reveals the presence of LMF1 at contact sites

Using standard IFA and fluorescence microscopy, LMF1’s staining pattern follows the length of the mitochondrion, as previously reported (Fig. 3A, (Jacobs et al., 2020)). In addition, we can detect patches of LMF1 in close proximity to the pellicle of the parasites (Fig. 3A, inset). With the advent of ultrastructure expansion microscopy (U-ExM), we revisited the localization of LMF1 to increase the level of detail and observe the distribution of the protein within the cell with higher resolution. Parasites expressing both LMF1(HA) and IMC10(Myc) were expanded in a water expansible acrylate gel, and the gels were stained with anti-HA and anti-Myc antibodies. We also stained the gels with *N*-hydroxysuccinamide ester (NHS-Ester), which binds to all primary amines of proteins and can reveal cell structures with a high level of detail (Dos Santos Pacheco et al., 2021). NHS staining allows us to visualize parasite structures such as the conoid, rhoptries, and the nucleus. Importantly, NHS also stained the mitochondrion, which allows us to track its shape without the use of an antibody (Fig 3B). In expanded parasites, IMC10 is uniformly distributed along the parasite’s periphery (Fig. 3B, magenta), and LMF1 follows the shape of the mitochondrion (Fig 3B, yellow and NHS-Esther). Additionally, we can observe that LMF1 is distributed to both sides of the outer mitochondrial membrane (Fig. 3C). Interestingly, we detect LMF1 dots in regions where the mitochondrion is near the pellicle, and in some of these contact regions, there is proximity between LMF1 and IMC10 staining (Fig. 3C). Drawing a line in an area where we see the mitochondrion, LMF1, and IMC10 near each other, we can determine the position of the peak intensity for each of the signals (Fig. 3C). This analysis shows that the peak intensity for LMF1 is near that of IMC10, suggesting proximity (Fig. 3C). We measured the distance between the peak intensity of LMF1 and IMC10 in 30 different parasites and calculated an average distance of 25±6.1 nm, which is consistent with a membrane contact site (MCS) (Jing et al., 2020).

**Figure 3.**
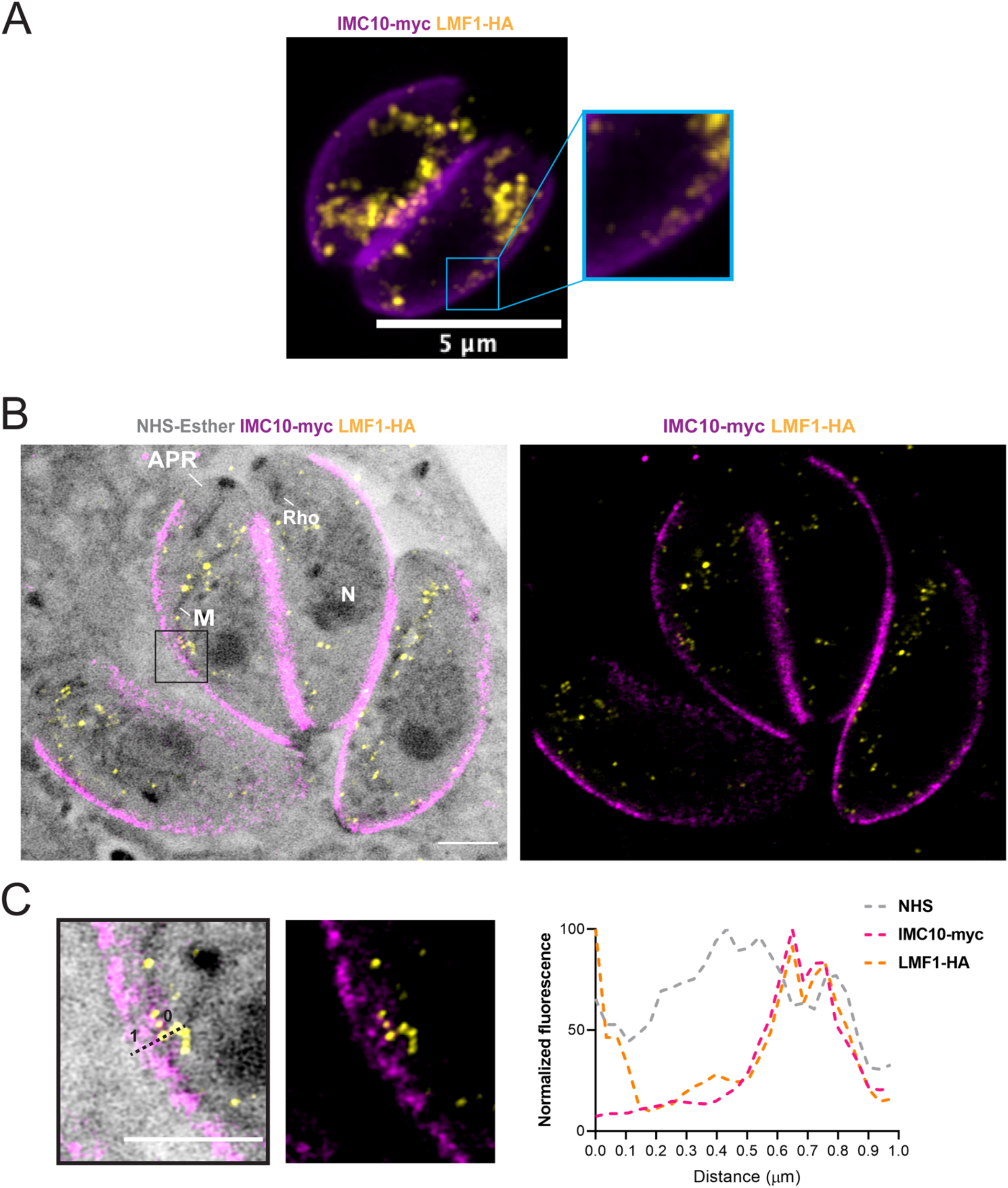
Expansion Microscopy shows colocalization between IMC10 and LMF1. A) IFA of intracellular parasites stained with anti-HA (yellow) to detect LMF1 and anti-myc (magenta) to detect IMC. Box highlights a portion of the cell where the two signals are adjacent. B) Ultrastructure Expansion Microscopy (U-ExM) of intracellular parasites stained for LMF1-HA (yellow) and IMC10-Myc (magenta). On the left is the fluorescence signal showing the distribution of both proteins in the expanded parasites. On the right is an overlay of that image with the signal for NHS-ester, a total protein density marker. NHS staining allows for the visualization of structures such as the apical polar ring (APR), mitochondrion (M), rhoptries (Rho), and nucleus (N). C) Enlarged images from the boxed area in C showing IMF1 and IMC10 in proximity to each other. Line marks a region of proximity between signals LMF1 and IMC10 signals, which was used to map fluorescence intensity for each signal. The graph shows the normalized fluorescence intensity (%) corresponding along the 1 µm line in the image on the left. Gray dotted line = NHS signal, pink dotted line = IMC10 signal and yellow dotted line = LMF1 signal. All scale bars in this panel = 5 µm. The fluorescence intensity was calculated using ZEN Blue Software.

### LMF1 associates with the parasite pellicle

The parasite’s pellicle is a detergent-resistant structure that can be isolated intact away from the rest of the parasite using deoxycholate (DOC). Accordingly, we isolated the parasite pellicle using 1% DOC and analyzed it by IFA and Western blot to confirm the presence of LMF1. IFA of the isolated pellicles shows staining for IMC10 and, importantly, also for LMF1 (Fig. 4A). Western blot analysis of the isolated pellicles reveals the presence of IMC10 and LMF1 in the pellicle fraction, but not of the inner mitochondrial membrane-localized ATP synthase beta subunit (Fig. 4B). These results using organelle extraction confirm the interaction of LMF1 with the pellicle, which is consistent with a role for LMF1 in mediating the contacts between the mitochondrion and the IMC.

**Figure 4.**
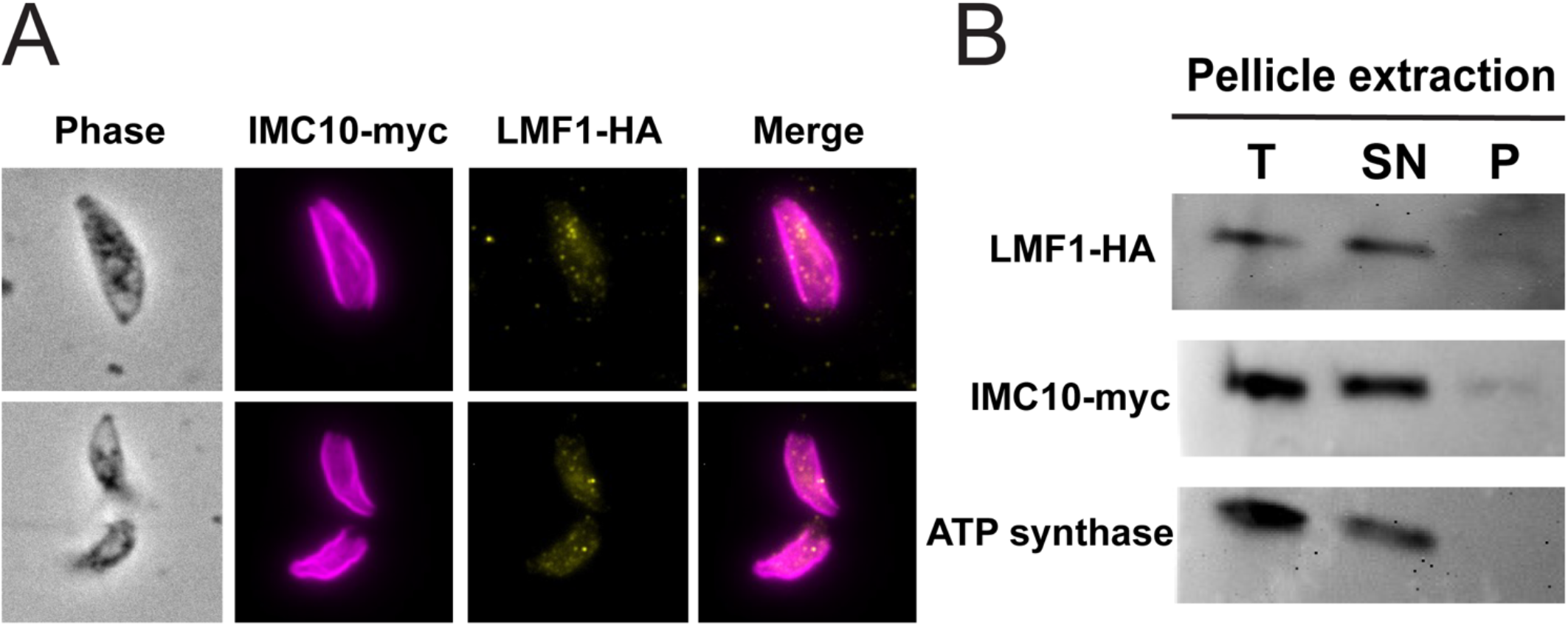
Localization of LMF1 in isolated pellicles. Pellicles were extracted from intracellular parasites with deoxycholate (DOC) and used for IFA (A) and Western Blots (B). A) IFA of DOC extracted pellicles showing IMC10 (magenta) and LMF1 (yellow). B) Representative western blots from total parasite extract (T) and both the supernatant (SN) and pellet (P) from the DOC extracted fraction. Western blots were probed for LMF1 (anti-HA), IMC10 (anti-Myc), and the ATP synthase subunit B. Only IMC10 and LMF1 are detected in the pellet fraction, which contains the parasite pellicle.

### IMC10 is not essential for *in vitro* propagation but is critical for mitochondrial morphology

Based on a whole-genome CRISPR selection IMC10 is predicted to be fitness conferring for tachyzoites in tissue culture (fitness score -4.01) and likely to be essential (Sidik et al., 2018). Accordingly, we generated a conditional knockdown strain by replacing the endogenous *IMC10* promoter with a tetracycline repressible one (Fig. 5A). We confirmed the insertion by PCR (Fig. 5B) and showed that the addition of the tetracycline analog ATc results in a significant reduction of *IMC10* mRNA levels in the TATI-*IMC10* but not in the parental strain (Fig. 5C).

**Fig 5.**
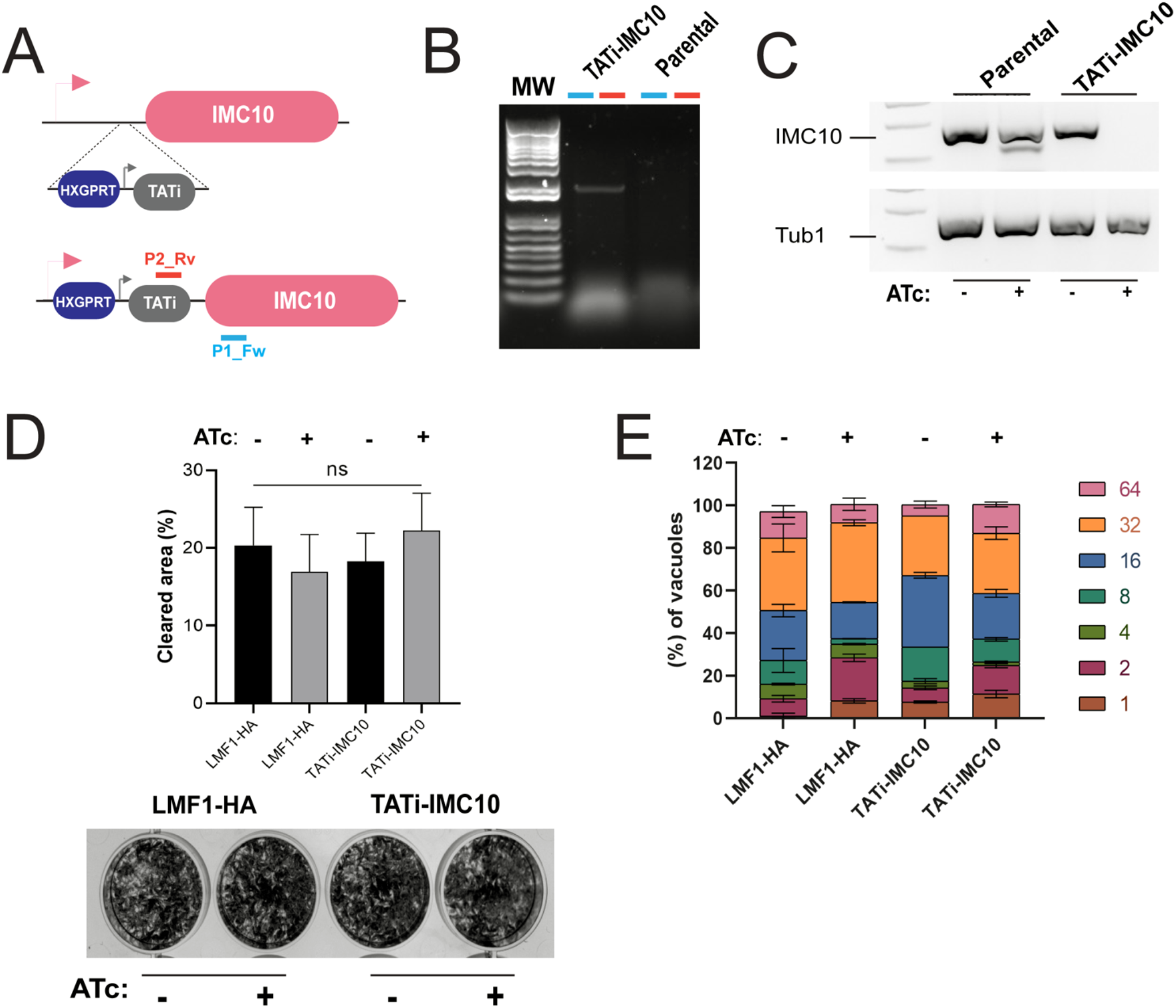
Conditional knockdown of IMC10 does not affect parasite propagation *in vitro*. A) Schematic representation of replacement of the endogenous IMC10 promoter for the TATi promoter cassette, which allows for repression of IMC10 by addition of the tetracycline analog ATc. P2_Rv and P1_Fw indicate the positions of the primers used to confirm the promoter replacement. B) PCR confirmation of promoter replacement using the primers depicted in A, which are expected to amplify a 2,200 base pairs amplicon in the TATi strain but not the parental. Primers are represented in A. C) Representative PCR reaction using cDNA produced from parasites of the parental strain and the TATi-*IMC10* strain grown with and without ATc for 24 hours. PCR reaction was done using specific primers for *IMC10* and tubulin, amplifying approximately 150 bp for each gene. D) The average number of plaques per well for either parental and knockdown cell lines grown with and without ATc after 5 days incubation period. Plaque assays were done in biological replicates (n=5), with error bars representing standard deviation (SD). ns = not significative (P> 0.05). E) Doubling assay. Parasites were allowed to invade HFFs for 1 h, and the cultures were fixed after 48 hours post-infection. The percentage of vacuoles containing 2, 4, 8, 16, 32, or 64 parasites was calculated for each condition. Doubling assays were performed in biological replicates (n=2). Error bars represent standard deviation (SD).

To assess if the knockdown would interfere with the replication of these parasites, we performed a plaque assay in which the parasites were incubated for five days with or without anhydrotetracycline (ATc). The plaque area cleared after five days was similar among the conditions and strains tested, showing that in the absence of IMC10, the parasites are still able to complete a full intracellular cycle. To confirm the lack of a growth phenotype, we performed a doubling assay with the TATi-*IMC10* and parental strains in the presence or absence of ATc. After 48h, we counted the number of parasites within the vacuoles and tabulated the percentage of vacuoles with a specific number of parasites across strains and conditions. As indicated by the plaque assay, no proliferation phenotype was observed with all conditions having the same distribution of vacuole sizes (Fig. 5E). In sum, these results indicate that contrary to what was suggested by the low fitness score, IMC10 is not essential for parasite propagation in tissue culture.

Given the interaction between LMF1 and IMC10 and the role of LMF1 in mitochondrial morphology, we determined whether the lack of IMC10 affected the normal mitochondrial dynamics. Normally intracellular parasites present mostly lasso-shaped mitochondrion, while extracellular ones present both sperm-like and collapsed mitochondrion (Jacobs et al., 2020; Ovciarikova et al., 2017). Lack of LMF1 leads to most intracellular parasites having either sperm-like or collapsed mitochondrion (Jacobs et al., 2020). Accordingly, we monitored the morphology of mitochondrion in TATi-*IMC10* parasites grown in the presence and absence of ATc (Fig. 5A). In addition, to focus on non-dividing parasites with an intact IMC, we also stained for IMC3 (Fig. 5A)(Gubbels et al., 2004). After 24 h post-infection, parasites without ATc showed a higher percentage of lasso (62.16±6.75%) in comparison to parasites in the presence of ATc (25.23±10.86%) (Fig. 5B). In addition, parasites under ATc treatment showed an increase in sperm-like shape (41.1±7.37% vs. 36.33±4.77% for parental) and collapsed mitochondrion (30.01±2.58% vs. 1.5±1.08% for parental). This phenotype is also observed after 48 h infection in the presence of ATc, with only 14%±2.01 of parasites showing lasso mitochondrion in contrast to 61.01±2% of parental (Fig. 5C).

We used U-ExM and NHS-ester staining to analyze the cell structure of parasites lacking IMC10 (Fig. 6). As with IFA, ExM reveals clear disruption of mitochondrial morphology when the TATi-*IMC10* parasites are exposed to ATc (Fig. 6). Figure 6A shows a vacuole of the knockdown strain grown in ATc showing all three morphologies: lasso, sperm-like, and collapsed (Fig. 6A). Interestingly, we note that while the mitochondrion appears lasso-like in some parasites, it is not continuous, and exhibits breaks along its length (Fig. 6A). As the expansion per se does not alter the mitochondrial morphology or its positioning, it was possible to observe regions of apposition of the mitochondrion and the parasite pellicle and to measure the distance between the two structures. When we induced the knockdown, it was difficult to find fields of view with a mitochondrion-pellicle contact site. To quantitate this observation, we selected 30 parasites that still exhibited a lasso in each condition and measured the distance between both structures. Based on our measurements, untreated parasites showed a distancing of 24.67±7.78 nm, closely related to what was previously described (26.23±12.02 nm (Ovciarikova et al., 2017)). There was a significant shift in the distance from the mitochondrion to the pellicle in the treated knockdown strain, with an average of 36.7±16.6 nm (Fig. 7B).

**Figure 6.**
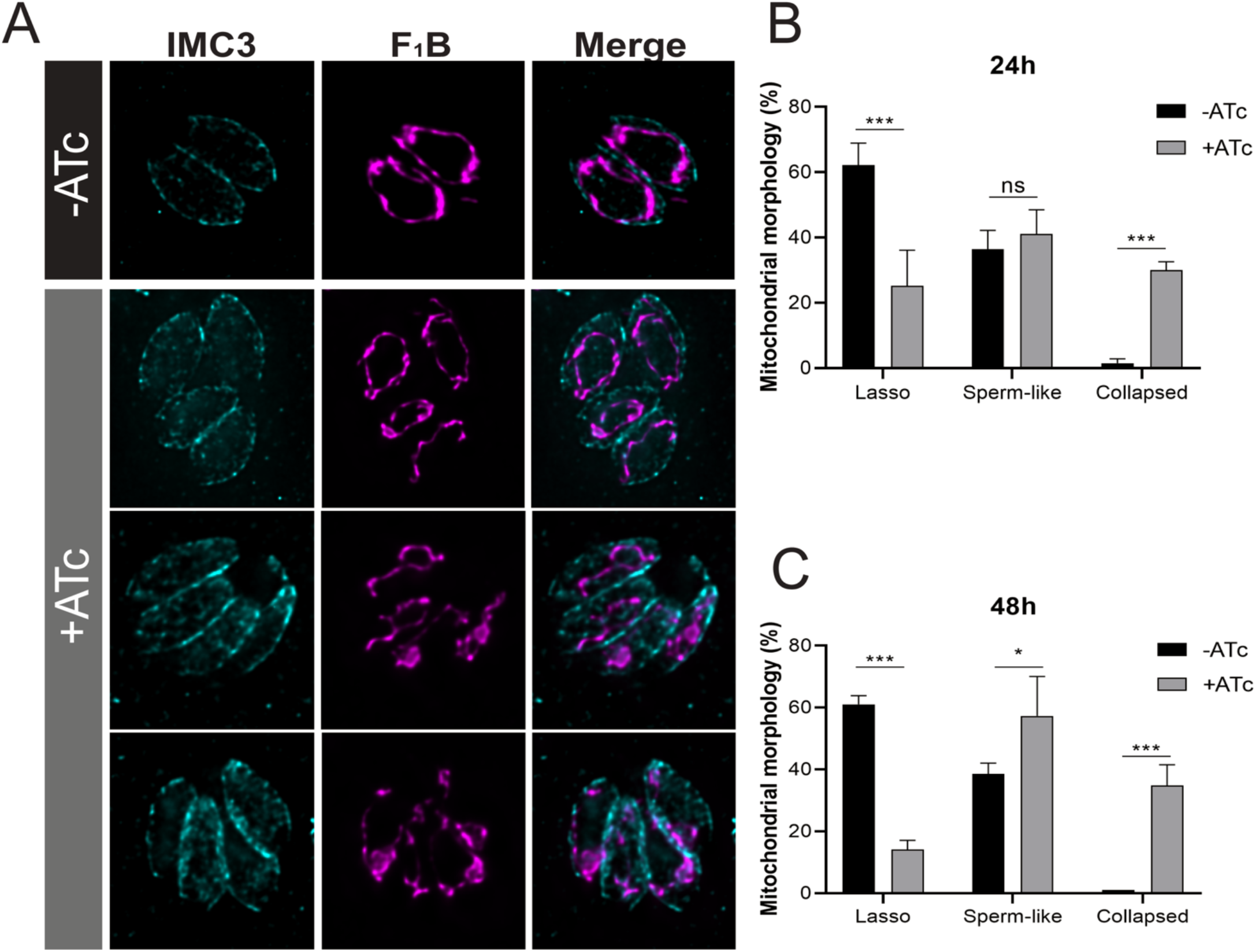
IMC10 knockdown disrupts mitochondrial morphology. A) Intracellular parasites of the TATi-IMC10 strain were grown without (-) or with (+) ATc to regulate IMC10 expression. Parasites were stained for IMC3 (cyan) and F_1_B-ATPase (magenta). Scale bar, 5 µm. B) and C) Percentage of parasites with each of the three different morphologies for parasites grown in the absence and presence of ATc after 24 and 48h. Data are an average of three replicates; at least 150 non-dividing vacuoles with intact IMC per sample were counted. *** p value < 0.001; ** p value < 0.01; * p value < 0.05. For a p value > 0.05, we consider the differences to be not significant (ns). Errors bars mean SD.

**Figure 7.**
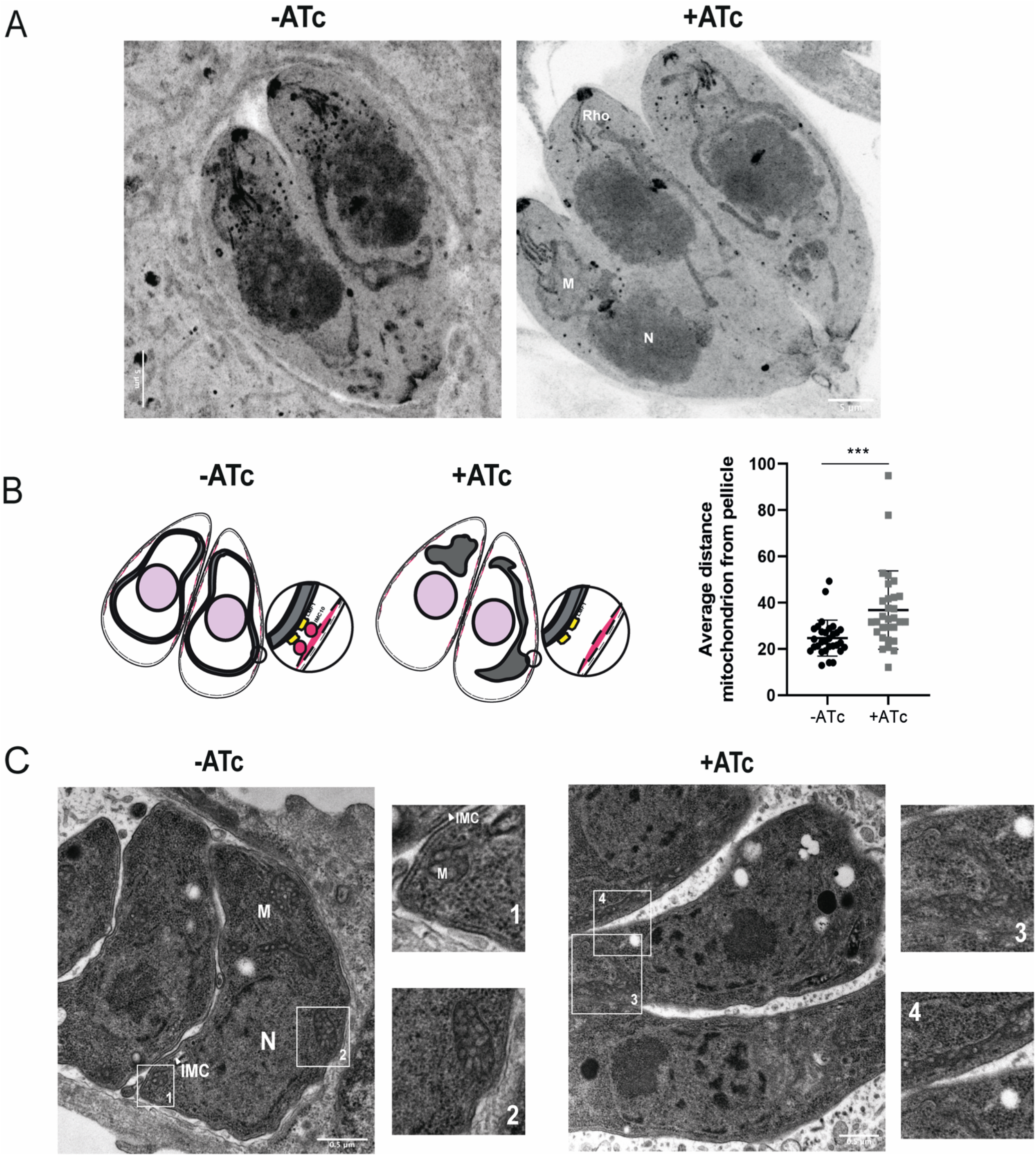
TATi-IMC10 cell lines show defects in mitochondrion position. A) Representative figure of parasites in the presence or absence (control) of ATc visualized by ExM. Parasites were expanded and stained with NHS-ester (protein density marker) to highlight cellular structures, such as the rhoptries (Rho), nucleus (N), and mitochondrion (M). Scale bar = 5 µm. B) Average distance from the pellicle to the mitochondrion was calculated in expanded parasite by measuring the distance between both organelles in their closest point. A total of 30 parasites were counted in two biological replicates (n=2). (***) p < 0.001. Error bars mean SD. C) Representative transmission electron microscopy (TEM) images of induced and non-induced cell lines 24h post-infection. Scale bars = 500 nm. Insets 1 to 4 show detailed structures, including the inner membrane complex (IMC), the nucleus (N), and the mitochondrion (M). Scale bars represent 500 nm for full pictures and 100 nm for the insets.

It is interesting to note that sometimes sperm-like mitochondrion is accumulated very close to the IMC, although it is not possible to know if this is the result of active tethering or if it is just a random effect. By U-ExM we can observe that the sperm-like mitochondrion is extending along the cell body towards the apical end of the parasite. No differences were found in cristae composition and distribution, and parasites lacking IMC10 do not show any significant structural differences in apicoplast and endoplasmic reticulum (Supplemental Figure S2). Transmission electron microscopy (EM) images also show clear differences in mitochondrial positioning after IMC10 ablation (Fig. 7C). In non-treated cells, it is possible to visualize patches of the mitochondrion in contact with the IMC (Fig. 7B boxes 1 and 2). Upon IMC10 knockdown, most parasites exhibit sperm like mitochondrion and/or collapsed ones. Interestingly, it was difficult to find slices showing both organelles in good resolution in the parasites lacking IMC10. In the +ATc cells, most of the mitochondrion is found collapsed onto the IMC, which makes it difficult to measure the distance between those organelles (Figure 7C insets 3 and 4). In contrast, we could measure the average distance from the mitochondrion to the pellicle in -ATc cells. Among 40 different TEM images, we observed 18 with visible mitochondrion and IMC, with an average distance between them of 58.28±21.86 nm. Together, these results suggest that the presence of IMC10 is critical for the mitochondrial morphology in intracellular parasites and likely plays a role in tethering the parasite’s mitochondrion to the pellicle.

### IMC10 iKD affect cell division and mitochondrial inheritance

As noted above, lasso-shaped mitochondrion in the absence of IMC10, while present in some parasites, did not appear contiguous as normal (Fig. 7A, arrow). Given that, we counted 150 vacuoles where the mitochondrion was in lasso shape in the + and -ATc parasites. In knockdown parasites grown without ATc, only 2.34±0.56 % of parasites exhibited a broken lasso. By contrast, after 24 h in ATc, the percentage of broken lasso-shaped mitochondrion increased to 32.3±6.5% (Fig 8A). We observe the same phenotype after 48 h in ATc (Figure 8A).

**Figure 8.**
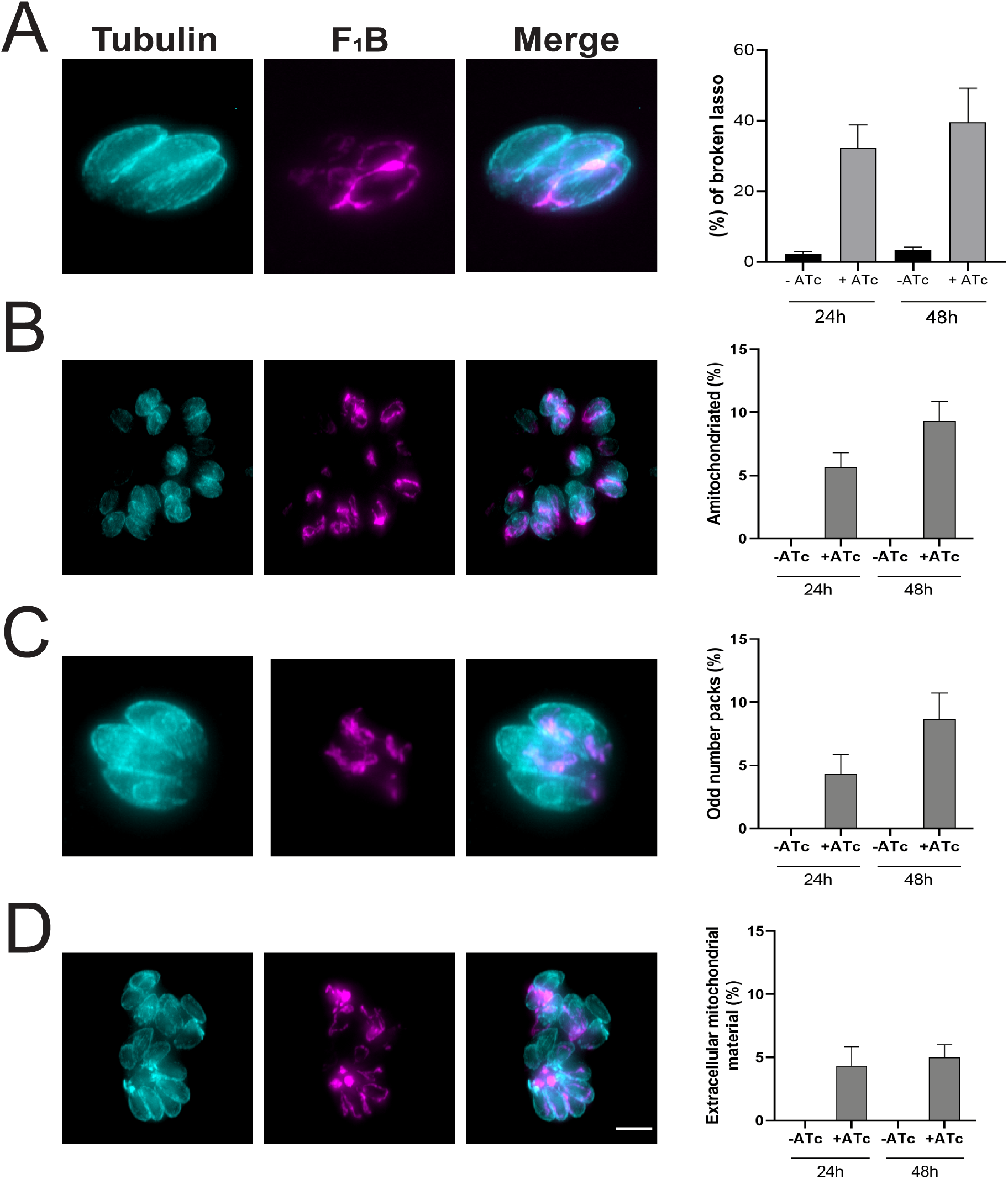
IMC10 knockdown exhibit another mitochondrial distribution phenotypes. IFA of knockdown parasites stained for Acetylated tubulin (cyan) and F_1_B-ATPase (magenta) showing aberrant phenotypes. A) Broken lasso. B) Amitochondriate parasites. C) Odd number of parasites in a vacuole and D) Accumulation of mitochondrion material outside of the cells within the same vacuole. Scale bar = 5 µm. All graphs represent the percentage of vacuoles with the related phenotype. At least 150 vacuoles per sample were inspected. For all graphs, n = 3. Error bars show SD.

During inspections of IFAs of parasites lacking IMC10, we detected numerous other phenotypes, probably related to defects during cell division upon IMC10 knockdown. Specifically, we detect parasites without a mitochondrion (amitochondriate) (Fig 8A), vacuoles with an unusual number of parasites (Fig 8C), and accumulation of mitochondrial material outside of the parasites (Fig 8D, arrowhead). All these phenotypes were quantitated, and we observed statistically significant differences between the parasites grown without and with ATc for either 24 or 48 hours (Figs 8B-D). Interestingly, all these phenotypes were observed in the LMF1 knockout parasite strain.

Normally during endodyogeny, two daughter cells form within a mother parasite. Nonetheless, events of more than two daughter cell budding can occur, albeit very rarely for wild-type parasites (Hu et al., 2002). As we observed the appearance of an abnormal number of parasites within the same vacuole, we decided to look if this phenotype is related to a defect during the division process. To monitor daughter cells, we stained parasites grown with and without ATc with IMC6. We considered any parasite with either one or more than two daughter cells as undergoing abnormal division. We observed that after 48h in the presence of ATc, the number of cells showing abnormal division was 12.3±0.8% within the population, which is significantly higher than the 1.5±0.8% observed in the absence of ATc (Fig 9A). Another characteristic observed upon IMC10 ablation was that parasites within the same vacuole were dividing asynchronously. Among all vacuoles with dividing parasites in the ATc grown parasites, 9.7±2% had parasites in different time points of division. By contrast, only 2.1±0.32% of those in the -ATc culture exhibited this phenotype. This result points out that IMC10 is important for cell division, even though the protein is not essential for the parasite’s lytic cycle.

**Figure 9.**
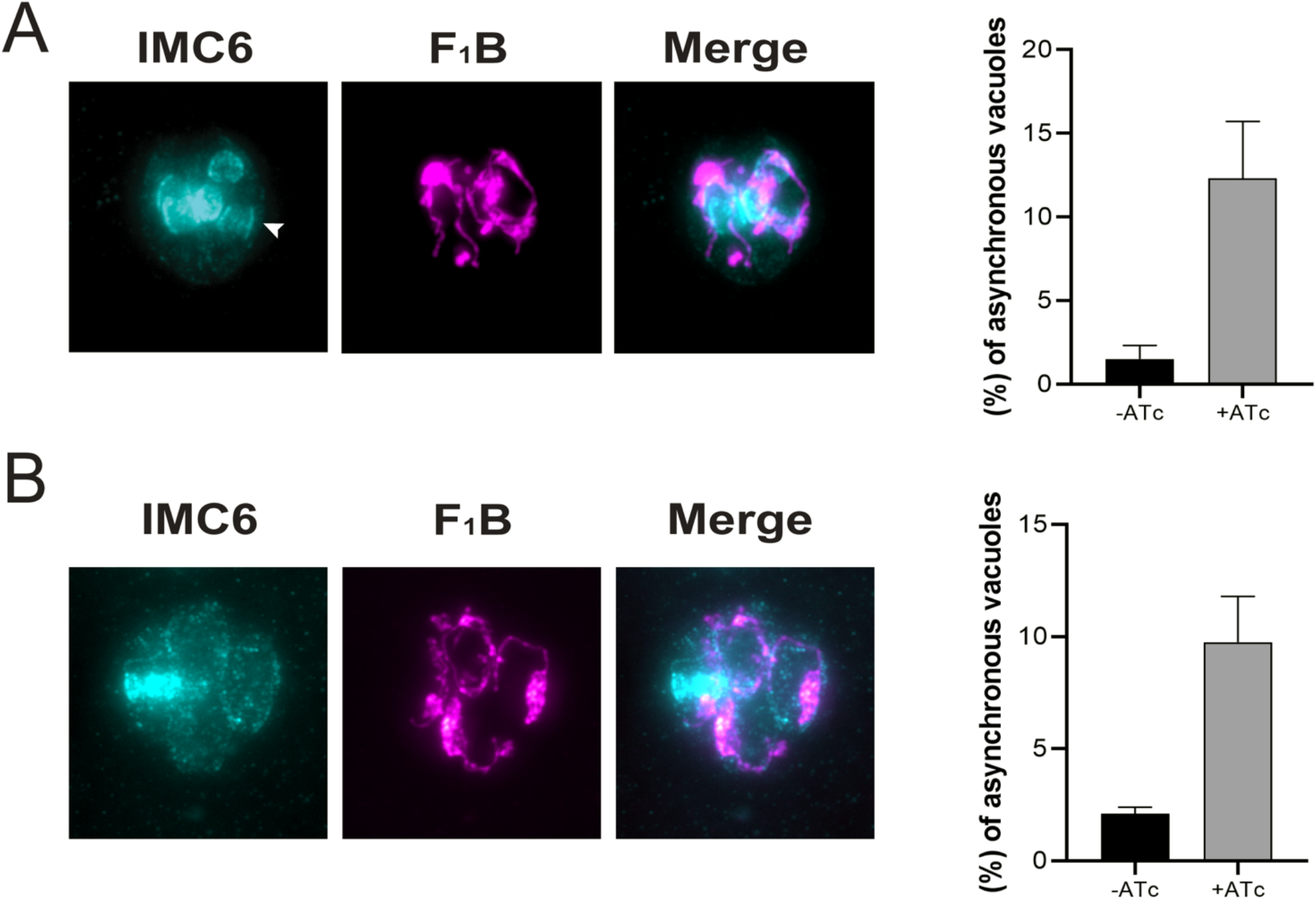
IMC10 knockdown cell lines exhibit division-related phenotypes. IFA of knockdown parasites stained for IMC6 (cyan) and F_1_B-ATPase (magenta) showing aberrant phenotypes. A) Aberrant number of budding cells within the same mother. B) Asynchronous vacuoles in which parasites are in different stages of division. Scale bar = 5 µm. All graphs represent the percentage of vacuoles with the related phenotype. At least 150 vacuoles per sample were inspected. For all graphs, n = 3. Error bars means SD.

### IMC10-LMF1 interaction promotes mitochondrial distribution during endodyogeny

Based on our results, IMC10 is important for mitochondrial distribution among daughter cells during division. As was mentioned before, mitochondrial distribution is also affected in cells lacking LMF1 (Jacobs et al., 2020). Accordingly, we examined the localization dynamics of these proteins during mitochondrial inheritance during cell division using U-ExM. We imaged the dual-tagged cell line LMF1(HA)/IMC10(Myc) at different time points of cell division and followed the distribution of the mitochondrion based on the NHS staining (Fig.10). It was previously observed that the mitochondrion is one of the last organelles to enter the daughter cells during endodyogeny (Nishi et al., 2008). During interphase, we can observe parasites presenting a full lasso, which appears to have contact with the parasite’s pellicle. As cell division progress, the lasso-shaped mitochondrion is opened, and it starts to move along the mother cell during early and mid-budding. Interestingly, the basal body of these stages is larger, and it is not possible to see any mitochondrion inside of the daughter cells. In late budding it is possible to observe that as the daughter cells grow, the basal complex tightens, and the mitochondrion branches start entering the daughter cells. The mitochondrion branches appear to be in close proximity to the daughter parasites’ pellicle (Fig. 10A and B).

**Figure 10.**
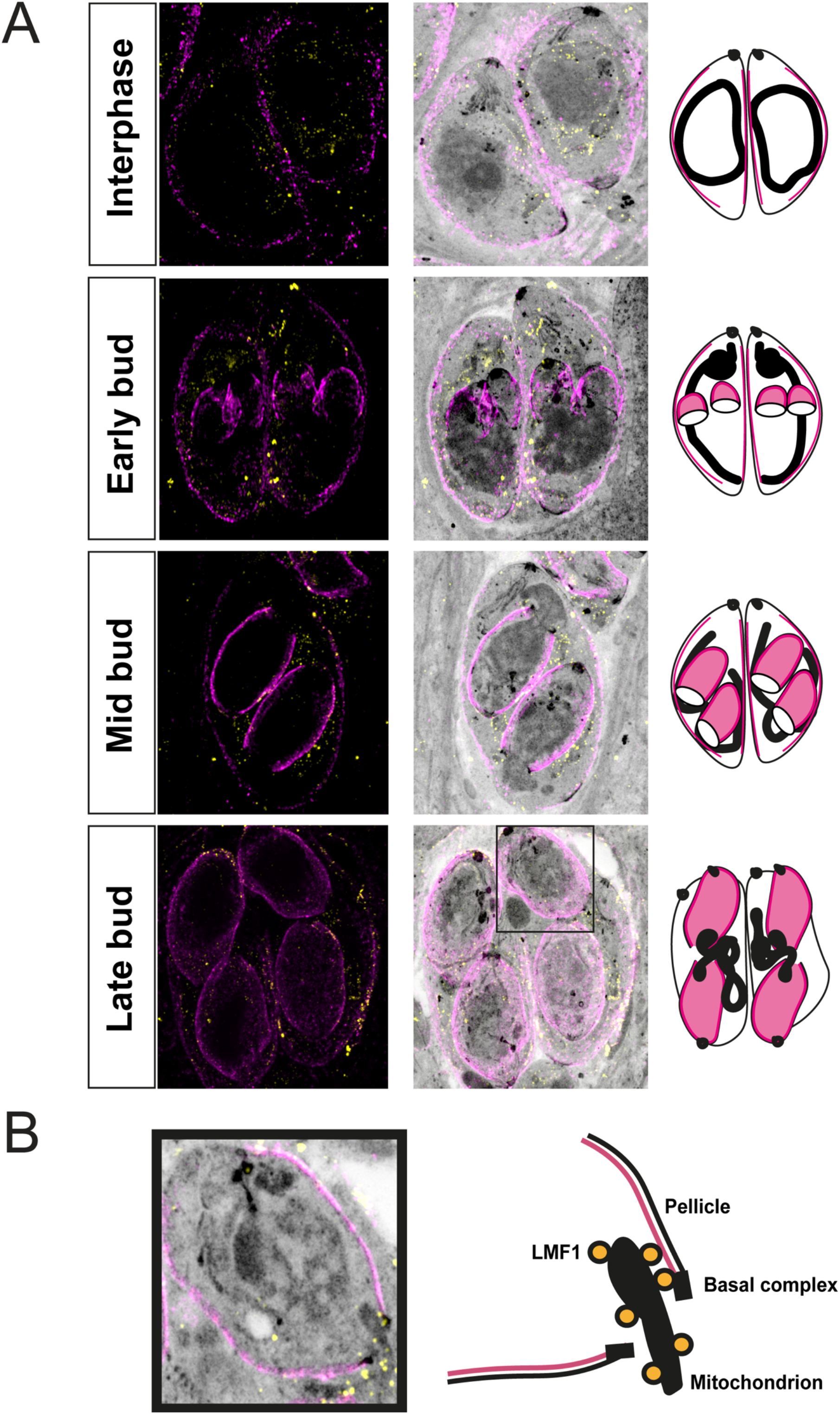
LMF1 and IMC10 interact during mitochondrial distribution. A) Ultrastructure Expansion Microscopy (U-ExM) of intracellular parasites stained for LMF1-HA (yellow) and IMC10-Myc (magenta). On the left is the fluorescence signal showing the distribution of both proteins in the expanded parasites followed by an overlay of that image with the signal for NHS-ester. On the right is a representation of the observed pattern of localization. The black square indicates the late budding cell used in B. Scale bar = 5 μm. B) Left: detail of a late-stage division cell with an emerging mitochondrion branch. Right: Scheme depicting the relative localization of the proteins, including structures such as the basal complex, mitochondrion, LMF1 (yellow circles), and pellicle.

At the same time, it is possible to observe that the proximity between LMF1 and IMC10 occurs during mitochondrion inheritance (Fig. 10B). Together, these results suggest that the MCS between the mitochondrion and the pellicle in *Toxoplasma* happens during cell division, and it is important for organelle division and distribution to the daughter cells, explaining the mitochondrial distribution phenotypes observed with a knockout of either LMF1 or IMC10.

## DISCUSSION

Membrane contact sites (MCS) are defined as regions where membranes from two compartments are tethered in close apposition (∼30 nm) and in which specific proteins and/or lipids are enriched (Eisenberg-Bord et al., 2016; Prinz, 2014). These contact sites are important for several physiological processes, including ion (Raturi et al., 2016) and lipid exchange (Aaltonen et al., 2022). In *Toxoplasma,* contact between organelles has been described between the mitochondrion and the apicoplast (Nishi et al., 2008), the endoplasmic reticulum (Mallo et al., 2021), the IMC (Jacobs et al., 2020; Ovciarikova et al., 2017) and between the ER and the apicoplast (Tomova et al., 2009). Nonetheless, neither the function nor the tethering proteins have been determined for these potential membrane contact sites. The one exception is the contact between the parasite mitochondrion and the pellicle. We previously described the identification and characterization of a coccidian-specific protein, LMF1, that localizes at the mitochondrion outer membrane and that is essential for positioning the mitochondrion to the periphery of the parasite (Jacobs et al., 2020). In the absence of LMF1, the mitochondrion loses its typical lasso shape in intracellular parasites and collapses to one of either end of the parasite. Accordingly, we hypothesized that LMF1 is part of a tethering complex that links the OMM to the parasite pellicle.

In this study, we focused on the proteins that collaborate with LMF1 in mitochondrion shaping and distribution. We confirmed three putative interactors first identified through a yeast two-hybrid screen: IMC10, a hypothetical EF-hand protein (TGGT1_246720), and ATPase-GC. TGGT1_246720 is present in the conoid of the mother cell, but it is also present in what appears to be the IMC of the daughter cells. ATPase-GC is an integral membrane protein localized towards the apical end of the parasite that is involved in Ca^2+^, phosphatidic acid (PA) signaling during egress, motility, and microneme secretion (Bisio et al., 2019; Brown and Sibley, 2018; Yang et al., 2019b). While TGGT1_246720 and ATPase-GC have been previously characterized, those studies did not investigate the shape of the mitochondrion in their absence. Given the rapid transition of the mitochondrion from a lasso to a collapsed morphology as the parasites exit the host cell, it is plausible that ATPase-GC and other signaling proteins that regulate egress are involved in regulating mitochondrial morphology. Similarly, the presence of TGGT1_246720 in daughter cells could suggest that its interaction with LMF1 is related to mitochondrial inheritance. Further work is needed to understand the role of these two proteins in the mitochondrial dynamics in *Toxoplasma*.

Due to the likely contact between the OMM, where LMF1 is localized, and the parasite pellicle, we were particularly intrigued by those interactors that are known to be part of the IMC: ILP1 and IMC10. The inner membrane complex is part of the parasite pellicle, and it is composed of flattened sacs termed alveoli, supported by the subpellicular network (Mann and Beckers, 2001) on the cytoplasmic face and interacts with the parasite’s microtubule cytoskeleton (reviewed by (Harding and Meissner, 2014) (Harding et al., 2019)). Attempts to tag ILP1 in the LMF1(HA) expressing strain failed, so we could not confirm the interaction. Nonetheless, disruption of ILP1 has been reported to affect the mitochondrion (Chen et al., 2015), although, given the pleiotropic effects of ILP1 knockdown, it is unclear whether this effect is specific. Nonetheless, we were able to confirm the interaction between LMF1 and IMC10 with various complementary approaches. IMC10 is an alveolin containing protein localized to the parasite’s pellicle (Anderson-White et al., 2011) and its expression is upregulated during cell division (Behnke et al., 2010). Surprisingly, the knockdown of IMC10 did not affect fitness, suggesting that it is not essential for propagation in tissue culture. This contrasts with what would have been expected based on the negative fitness score from a CRISPR high-throughput screen (Sidik et al., 2016). It is possible that under our knockdown conditions, some protein is still expressed, which is enough to maintain normal propagation. Interestingly, other IMC proteins with the same expression pattern as IMC10, such as IMC14 and 15, also showed not to be essential for parasite propagation in culture (Dubey et al., 2017). Regardless, knockdown of IMC10 significantly affected the mitochondrial morphology in intracellular parasites, phenocopying the effects of knocking out LMF1. The fact that lack of either LMF1 or IMC10 results in the same phenotypes and that we confirmed their interaction with three different approaches strongly suggest that these two proteins are part of a coccidian-specific tethering complex. A study recently reported that the parasite’s porin mediates the contacts between the mitochondrion and the ER. Knockdown of a mitochondrial porin leads to morphological defects in both the mitochondrion and the ER (Mallo et al., 2021). Thus, it is evident that contact and tethering to other structures of the parasite are central to the mitochondrion’s morphology.

In yeast, a protein called Num1 is the tether that mediates mitochondrial-cortex contacts sites and confers proper mitochondrial segregation during cell division (Kraft and Lackner, 2017). This protein is structurally composed of internal EF-hands, a coiled-coiled domain that binds to the mitochondria, and a Pleckstrin homology domain (PH domain) that binds to the plasma membrane lipids (Ping et al., 2016). In silico analysis shows that LMF1 has no lipid-binding domains, which reinforces the idea that protein-protein interactions mediate *Toxoplasma*’s mitochondrial dynamics. Our previous studies suggest that LMF1 associates with the OMM through an interaction between its C-terminal domain with Fis1 (Jacobs et al., 2020). In the LMF1 Y2H, the interaction region of IMC10 to LMF1 is a part of the IMC10 C-terminal, probably connecting to the N-terminal of LMF1. More studies are necessary to describe the minimal region that determines the interactions among these proteins. Regulation of mitochondrial division in related apicomplexans such as *Plasmodium* is still an open question. Recently it was reported that a Fis1 homolog in *Plasmodium* is dispensable for mitochondrial division. As knockout of PfFis1 showed no effect in either parasite growth or mitochondrial division, the authors hypothesized that other proteins are participating in this process (Maruthi et al., 2020). Interestingly, *Plasmodium* does not seem to encode an LMF1 homolog.

IMC10 knockdown affected not only mitochondrial morphology but also cell division. After induction, it was possible to observe asynchronous cell division, leading to an increase in the number of polyploid cells and vacuoles with an abnormal number of parasites. These division defects might also relate to the defects in mitochondrial distribution to the daughter cells, which result in amitochondriate parasites and excess extracellular mitochondrial material within the vacuoles. Importantly, lack of LMF1 also results in mitochondrial inheritance defects, suggesting that the LMF1/IMC10 complex plays a role during endodyogeny.

Mitochondrial division in *Toxoplasma* is a tightly regulated process within daughter cell budding (Nishi et al., 2008). The organelle is one of the last to enter the newly formed cells, and it is possible to observe branches of this organelle emerging and entering daughter cells in the late endodyogeny stages (Nishi et al., 2008; Verhoef et al., 2021). Using U-ExM, we could observe what appear to be LMF1-IMC10 complexes upon mitochondrial distribution to the daughter cells. This data strongly suggests that the formation of MCS is important for mitochondrial inheritance, given the fact that disturbing both IMC10 and LMF1 causes mitochondrial segregation defects and excessive accumulation of mitochondrial material in the residual body. The understanding of the proteins involved in mitochondrial division in apicomplexan parasites is still very limited (Verhoef et al., 2021; Voleman and Dolezal, 2019). Apicomplexan parasites do not appear to encode homologs of bacterial FtsZ and instead encode a set of dynamin-related proteins (Drp) (Morano and Dvorin, 2021). In *Toxoplasma*, DrpA is involved in apicoplast division (van Dooren et al., 2009), DrpB in secretory organelles biogenesis (Breinich et al., 2009) and DrpC play a role in vesicle transport (Heredero-Bermejo et al., 2019) and mitochondrial fission (Melatti et al., 2019), although the association of this protein with the mitochondrion is still unclear. Indeed, DrpC is intriguing because this protein lacks its GTPase Effector Domain (GED). The fact that there is not a direct involvement of a canonical dynamin-related protein and in the absence of Fis1 does not affect mitochondrial morphology in this parasite leads to questioning what are the other proteins mediating mitochondrial fission and inheritance in the parasite. Our identification of a novel and unique tethering complex that mediates mitochondrial contact with the pellicle provides a handle with which to study the morphodynamics of the mitochondrion of this important pathogenic parasite. Future studies of other components of this tethering complex, especially those involved in its regulation, will shed light on this important aspect of *Toxoplasma*’s biology and potentially reveal novel avenues for therapeutic interventions.

## MATERIAL AND METHODS

### Parasite culture and reagents

All the parasite strains were maintained via continued passage through human foreskin fibroblasts (HFFs) purchased from ATCC and cultured in Dulbecco’s modified Eagle’s medium (DMEM) high glucose, supplemented with 10% fetal calf serum (FCS), two mM L-glutamine, and 100 U penicillin/100μg streptomycin per mL. The cultures were maintained at 37°C and 5% CO_2_. Parasites used in this study were of the strain RH lacking hypoxanthine-xanthine-guanine phosphoribosyl transferase (HPT) and Ku80 (RHΔHPTΔku80) (ref).

### Phylogeny and domain prediction

Domain prediction was determined by the InterPro 87.0 (https://www.ebi.ac.uk/interpro/search/sequence/) tool using the full LMF1 amino acid sequence. To confirm the prediction of intrinsically disordered domains, we used the MobiDB (https://mobidb.bio.unipd.it) tool (Piovesan et al., 2021). Phylogeny was performed using the tBLASTn tool to compare amino acid sequences against the LMF1 protein. Homologs among apicomplexans and other organisms were confirmed by searches using the ToxoDB (https://toxodb.org/toxo/app) blast tool (Amos et al., 2022). For the phylogenetic tree, we used the OrthoMCL DB tool (https://orthomcl.org/orthomcl/app/) (Zdobnov et al., 2021). LMF1 orthologue group (OG6_176398) was used to detect homologs and determine their phyletic distribution. A cutoff of 1e-5 was used in this search. In total, 37 sequences were found for this phyletic group, with an average of 57.6% identity among all sequences. Out of the 37 sequences, we selected ten sequences for the phylogenetic tree. The tree was calculated using the tool Clustal Omega tool. Accessions for the sequences used in this work are: *Sarcocystis neurona* N3 (sneu|SN3_01200745), *Cystoisospora suis* strain Wien I (csui|CSUI_005550), *Besnoitia besnoiti* strain Bb-Ger1 (bbes|BESB_048460), *Neospora caninum* Liverpool (ncan|NCLIV_040070), *Toxoplasma gondii* GT1 (tggt|TGGT1_265180), *Toxoplasma* gondii ME49 (tgon|TGME49_265180), *Eimeria tenella* Houghton 2021 (etht|ETH2_1406700), *Cyclospora cayetanensis* strain CHN_HEN01 (ccay|cyc_05565), *Guillardia theta* (strain CCMP2712) (Cryptophyte) (gthe|L1IU32) and *Chromera velia* CCMP2878 (cvel|Cvel_23028).

### Yeast two-hybrid (Y2H)

Yeast two-hybrid (Y2H) screening was performed by Hybrigenics Services, S.A.S., Paris, France. The coding sequence for LMF1 (aa 1-452, XP_002368647.1) was PCR amplified and cloned into a pB66 as a C-terminal fusion with the Gal4 DNA-binding domain (Fromont-Racine et al., 1997). 46 million clones (5-fold the complexity of the library) were screened using a mating approach with YHGX13 (Y187 *ade2-101::loxP-kanMX-loxP, matα*) and CG1945 (*matα*) yeast strains as previously described (Fromont-Racine et al., 1997). 257 His (+) colonies were selected on a medium lacking tryptophan, leucine, and histidine. The prey fragments of the positive clones were amplified by PCR and sequenced at their 5’ and 3’ junctions. The resulting sequences were used to identify the corresponding interacting proteins in the GenBank database (NCBI) using a fully automated procedure. A confidence score (PBS, for Predicted Biological Score) was attributed to each interaction as previously described (Formstecher et al., 2005).

### Generation of endogenously dual-tagged cell lines

For the C-terminal endogenous tagging of LMF1 putative interactors, we introduced a cassette encoding a 3x-Myc tag directly upstream to the stop codon for the gene of interest. This cassette included the selectable marker HXGPRT and was amplified from the vector pLIC-3xmyc-HXGPRT (Huynh and Carruthers, 2009)with primers that included the homology regions of each gene to promote recombination. Insertion of the cassette was facilitated by CRISPR. For this purpose, we replaced the guide RNA in pSAG1-Cas9-GFP-pU6-sgKu80 (modified by (Blakely et al., 2020) from the original pSAG1-Cas9-GFP-UPRT (Shen et al., 2014)) for one targeting the *LFM1* locus using the Q5 site-directed mutagenesis kit (NEB). 1 µg of the cassette and 1 µg of Cas9 plasmid were transfected into the LMF1-HA cell line using the Lonza nucleofection system. Parasites were selected using mycophenolic acid (MPA), and independent clones were collected by serial dilution. All the primers used in this work are listed in Supplemental Table 2.

### Immunoprecipitation (IP) and co-immunoprecipitation (co-IP) assays

To confirm the results of the Y2H screening, we performed co-immunoprecipitations (co-IP) using the LMF1-HA cell line. Intracellular parasites from 10 T175 cultures were released by passing through a 21-gauge needle, spun down (1,000 x *g* 4° C), washed twice in cold PBS, and resuspended in Pierce co-immunoprecipitation lysis buffer (Thermo Fisher Scientific) with protease/phosphatase inhibitor cocktail (100X, Cell Signalling Technology). After 1 h of lysis at 4° C, the samples were sonicated three times for 20 s each time (20% frequency). After sonication, samples were pelleted, and the supernatant was incubated with anti-HA magnetic beads (Thermo Fisher Scientific). Samples were placed in a rocker for 2.5h before beads were washed once with Pierce co-IP lysis buffer and twice with PBS. Beads were resuspended in 8 M urea and sent for liquid chromatography coupled to tandem mass spectrometry (LC/MS-MS) analysis. Results were narrowed down to proteins that had at least four peptides in the LMF1-HA sample and none in control. To confirm the interaction between LMF1 and its putative interactors, we performed co-immunoprecipitation using the dually tagged cell lines. Intracellular parasites from 2 T175 cultures were syringe-released, and the samples were processed as described for the IP. In the end, the beads and total lysate were resuspended in 2X Laemmli sample buffer (Bio-Rad) supplemented with 5% 2-mercaptoethanol (Sigma-Aldrich) for western blot.

### Western blots

Parasite extracts were resuspended in 2× Laemmli sample buffer (Bio-Rad) with 5% 2- mercaptoethanol (Sigma-Aldrich). Samples were boiled for 5 min at 95°C before separation on a gradient 4 to 20% sodium dodecyl sulfate (SDS)-polyacrylamide gel (Bio-Rad). Samples were then transferred to nitrocellulose membrane using standard methods for semidry transfer (Bio-Rad). Membranes were probed with rabbit anti-HA (Cell Signaling Technologies), mouse anti-c-Myc (Cell Signaling Technologies), or mouse anti-F_1_B ATPase at a dilution of 1:5,000 overnight. Given the high molecular weight of ATPase-GC, we used 4-20% Tris-acetate SDS gels (Invitrogen) as performed in (Yang et al., 2019b). Membranes were then washed and probed with either goat anti-mouse horseradish peroxidase or goat anti-rabbit horseradish peroxidase (Sigma-Aldrich) at a dilution of 1:10,000 for 1 h (GE Healthcare). Proteins were detected using SuperSignal West Femto substrate (Thermo Fisher) and imaged using the FluorChem R system (Biotechne). All original Western blots are shown in Data Set S2 in the supplemental material.

### Duolink® Proximity Ligation Assay (PLA)

Dually tagged parasites syringe-released from host cells washed twice in cold PBS and fixed with 4% paraformaldehyde for 20min at room temperature. After fixation, cells were washed once in PBS and then seeded in poly-L-lysine (Sigma-Aldrich) coated glass coverslips. The cells were permeabilized using PBS + 0.25% Triton X-100 for 30 min at RT. DuoLink® assay (Sigma-Aldrich) was performed according to the manufacturer’s instructions with the following modifications: overnight blocking in a humidity chamber and five washes with 1 mL washing buffer per coverslip.

### Generation of IMC10 inducible knockdown strain

To generate the IMC10 inducible knockdown (iKD) strain, we introduced a cassette encoding a transactivator protein (TATi) and a tetracycline responsive element (TRE) upstream to the IMC10 start codon (Salamun et al., 2014). The cassette was amplified from the vector pT8TATi-HXGPRT-tetO7S (Salamun et al., 2014) with primers that included the homology regions corresponding to the upstream region of the *IMC10* gene. The Cas9 guide was made using the pSAG1-Cas9-GFP-pU6-sgKu80 (Blakely et al., 2020) as a template, and the sequences were introduced with the Q5 site-directed mutagenesis kit (NEB). Transfection of both the TATi cassette and the Cas9/guide RNA vector was performed as above. Correct integration of the TATi insert was validated by PCR. To confirm IMC10 knockdown, freshly lysed parasites were seeded in a confluent HFF monolayer in a T25 flask, with or without ATc. After 24h, parasites were syringe-released using a 21-gauge needle and washed twice with PBS (1 000 *g* x 4°C for 10 min). Total RNA was isolated using TRIzol following the manufactures instructions. RNA was treated with RNase-Free DNase and quantified by NanoDrop. A total of 500 ng of total RNA was used for cDNA synthesis using a SuperScript® III First-Strand kit (Invitrogen), following the manufacturer’s instructions. PCR reactions were performed using the cDNA to amplify a 150 bp product from IMC10 and Tubulin as a control. Primers are listed in Supplemental Table S2.

### Immunofluorescence assays

For all immunofluorescence assays (IFA), infected HFF monolayers were fixed with 3.5% paraformaldehyde, quenched with 100 mM glycine, and blocked with PBS containing 3% bovine albumin serum (BSA). Cells were permeabilized in PBS containing 3% BSA and 0.25% Triton X-100 (TX-100). Samples were then incubated with primary antibodies diluted in permeabilization solution for 1 h, washed five times with PBS, and incubated with the respective Alexa Fluor conjugated antibodies and 5μg/mL Hoechst (nuclear marker) in PBS for 1 h. The coverslips were washed five times with PBS. After washes, the coverslips were mounted in ProLong Diamond (Thermo Fisher Scientific). Image acquisition and processing were performed using either a Leica DMI6000 B microscope coupled with a LAS X 1.5.1.13187 software, a Nikon Eclipse 801 microscope with NIS-Elements AR 3.0, or a Zeiss LSM 800 AxioObserver microscope with an AiryScan detector using a ZEN Blue software (version 3.1). Images were processed and analyzed using FIJI ImageJ 64 Software. Primary antibodies used in this study are rabbit anti-HA (C29F4 Cell Signalling 1:1,000 dilution), mouse anti-Myc (9B11 Cell Signalling 1:1,000), rabbit anti-acetyl Tubulin (Lys40) (Millipore ABT241, 1:2,000), rat anti-IMC3 (1:2,000), rabbit anti-IMC6 (1:2,000), mouse anti-F1B ATPase (1:5,000), rabbit anti-ACP (1:5,000) and anti-SERCA (1:1,000). Secondary antibodies included Alexa Fluor 594- or Alexa Fluor 488-conjugated goat anti-rabbit and goat anti-mouse (Invitrogen), all used at 1:1,000.

### Ultrastructure Expansion Microscopy (U-ExM)

Ultrastructure Expansion Microscopy (U-ExM) was performed as described (Liffner and Absalon, 2021) with the following modification: the parasites were seeded in an HFF monolayer grown on glass coverslips in 24 well plates. Primary antibodies used are rabbit anti-HA (C29F4 Cell Signalling 1:100 dilution) mouse anti-Myc (9B11 Cell Signalling 1:500). Secondary antibodies used in this study are Alexa Fluor NHS 405, Alexa Fluor 594- or Alexa Fluor 488-conjugated goat anti-rabbit and goat anti-mouse (Invitrogen). Secondary antibodies were used at 1:1,000, except for the Alexa Fluor 405, which was used at 1:250.

### Pellicle extraction

Pellicle extraction was performed as previously described (Gilk et al., 2006) with some modifications. Briefly, 1 x 10^8^ parasites were resuspended in PBS containing 1% deoxycholate (DOC, v/v). After one cycle of sonication (20% frequency for 20 seconds on ice), the extract was centrifuged (15,000 x g at 4°C) for 30 minutes. Part of the extract was recovered for IFA, and the other part was boiled in Laemmli Buffer 2X (BioRad) supplemented with 5% 2-mercaptoethanol for western blot analysis.

### Phenotypic characterization of the IMC10 iKD strain

For the plaque assays, 500 freshly egressed parasites were seeded in a confluent HFF monolayer in 12-well plates, with or without ATc. After five days of incubation, cultures were fixed with methanol for 15 min and stained with crystal violet. Plaques were imaged using a Protein Simple imager, and the plaque area was calculated using the ColonyArea plugin (Guzman et al., 2014) on FIJI. The experiment was performed in five biological replicates, each with 3 technical replicates. Doubling assays were performed in 24-well plates as described before (Yang et al., 2019a). Percentage of vacuoles with a particular number of parasites (i.e., 2, 4, 8, 16, etc.) was tabulated for all strains, and conditions were monitored at 16, 24, and 40 h after infection. The experiment was performed in duplicate, each one containing three technical replicates.

For the mitochondrial morphology counts, HFFs infected with the iKD strain were grown with or without 0.5 μg/mL of Anhydrotetracycline (ATc) for 24 and 48 h. IFA was performed as described above using mouse anti-F_1_B ATPase and anti-rabbit IMC6 or rabbit anti-IMC3. Samples were examined blindly, and at least 150 non-dividing vacuoles with an intact IMC were inspected. Three mitochondrial morphological categories were quantitated: lasso, sperm-like, and collapsed (Jacobs et al., 2020; Ovciarikova et al., 2017). The other mitochondrial phenotypes accessed during the image analysis were counted and categorized as: broken lasso (Mallo et al., 2021), amitochondriate parasites, and extracellular mitochondrial material (Jacobs et al., 2020). The division phenotypes were quantitated using the same images by counting the number of synchronous and asynchronous vacuoles and the number of daughter cells in each dividing parasite. Synchronous vacuoles are those in which all parasites were in the same stage of division. At least 150 vacuoles were counted per condition. Experiments were performed in biological triplicates.

To calculate the distance from the mitochondrion to the pellicle in the expanded parasites, we inspected 30 parasites per condition (- and +ATc) in biological duplicates. The distance was measured based on the closest point between both organelles. Images were processed and analyzed using FIJI ImageJ 64 Software.

### Transmission Electron microscopy (TEM)

HFFs monolayers were infected with IMC10 iKD parasites for 24 h, in the presence (or not) of ATc. The cultures were washed 3x in PBS and fixed with 2% glutaraldehyde/2% paraformaldehyde in 0.1 M sodium phosphate buffer (pH = 7.4) for 1 h. After fixation, cells were harvested by scraping the monolayer and centrifuged at 1000 x g for 5 min. Cells were post-fixed for 95 min in the dark in 1% osmium tetroxide (OsO_4_) diluted in ultrapure water. After fixation, the cells were dehydrated in increasing concentrations of ethanol (50%-100%) at room temperature and embedded in EPON resin (Electron Microscopy Sciences). Ultrathin sections (70-80 nm) were obtained in a UCT Ultracut with FCS (Leica) and the sections visualized in a Tecnai Spirit OR (ThermoFisher) equipped with AMT CCD Camera (Advanced Microscopy Techniques). Images were acquired at the Electron Microscopy Core Facility at the Indiana University School of Medicine.

## ACKNOWLEDGMENTS

We want to thank Dr. David Sibley and Dr. Peter Bradley for sharing plasmids. We would like to thank Dr. Sabrina Absalon and Dr. Benjamin Liffner for their assistance with the expansion microscopy experiments and Dr. Irene Herdero Bermejo for assistance with the preparation of samples for transmission electron microscopy. This research was supported by the National Institute of Health grants R01AI123457, R01AI149766, RO1AI89808, and R21AI124067 to GA.

## SUPPLEMENTAL MATERIAL

**Supplemental Table S1.** LMF1 interactors identified by immunoprecipitation. Listed are proteins that had at least four peptides in the LMF1 IP and zero in control (parental parasites). LMF1 is in yellow and IMC10 in pink. Proteins that are known to localize to the pellicle are highlighted in grey. Included are the Gene ID, the gene annotation, and the number of peptides detected by mass spectrometry.

| Gene ID | Product Description | Total Peptides |
| --- | --- | --- |
| **TGGT1_265180** | **LMF1** | **53** |
| TGGT1_286580 | IMC17 | 40 |
| TGGT1_219320 | GAP50 | 38 |
| **TGGT1_230210** | **IMC10** | **30** |
| TGGT1_324600 | HSP20 | 26 |
| TGGT1_222220 | IMC7 | 23 |
| TGGT1_308860 | AC3 | 21 |
| TGGT1_232410 | TrxL1 | 20 |
| TGGT1_258410 | PhIL1 | 16 |
| TGGT1_248700 | IMC12 | 13 |
| TGGT1_241170 | hypothetical | 12 |
| TGGT1_252360 | ROP24 | 10 |
| TGGT1_219270 | GAPM2a | 9 |
| TGGT1_219310 | DnaK family protein (HSP70) | 9 |
| TGGT1_249970 | Acylated Pleckstrin-Homology (APH) | 7 |
| TGGT1_262050 | ROP39 | 6 |
| TGGT1_287500 | T complex chaperonin, putative | 4 |
| TGGT1_316540 | ISP3 | 4 |

**Supplemental Table S2.**
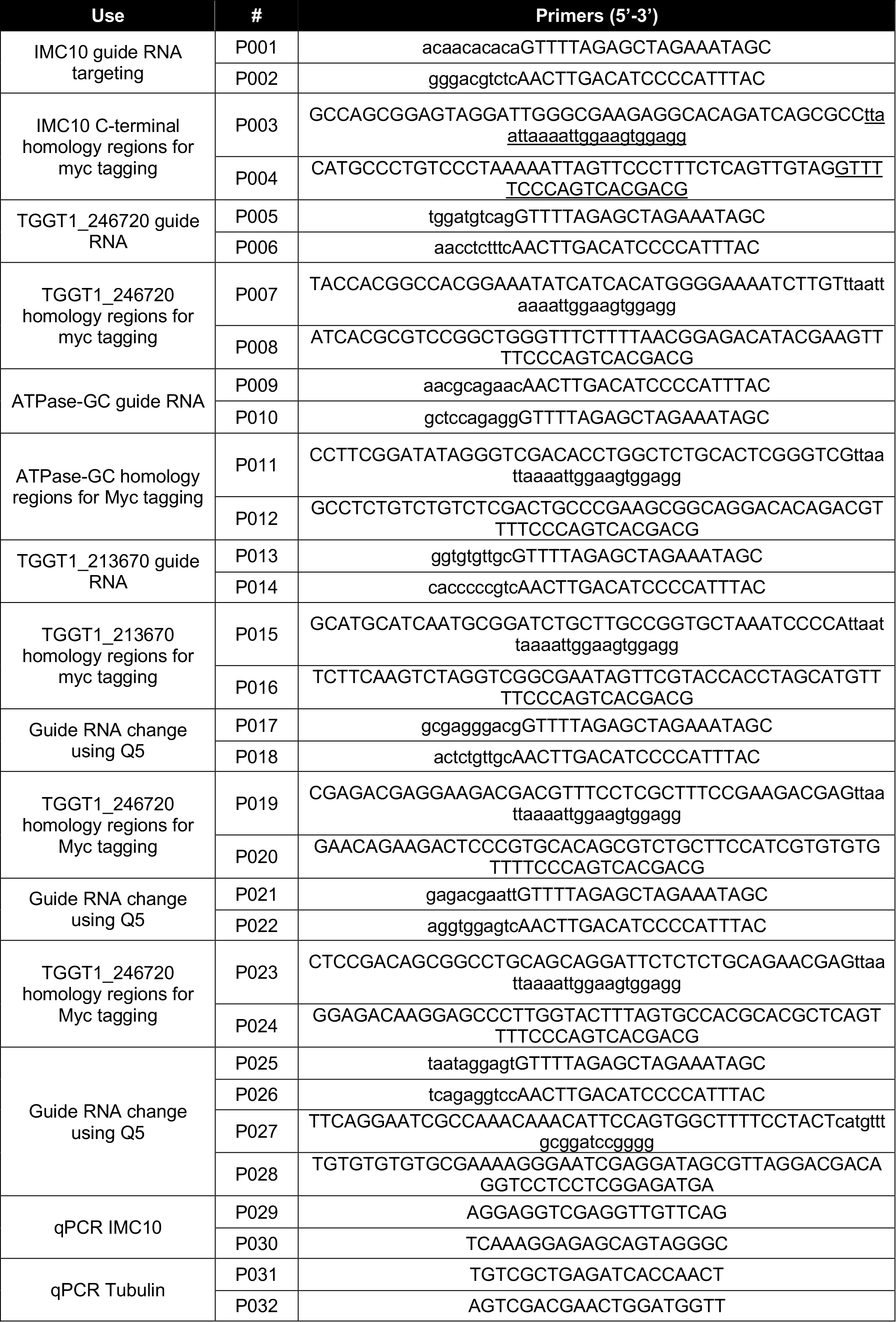
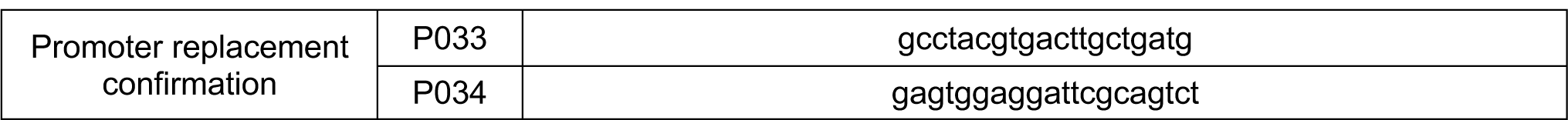
Sequences of primers used in this study. Small caps indicate overhangs with homology to the targeted region for recombination.

**Supplemental Figure S1.**
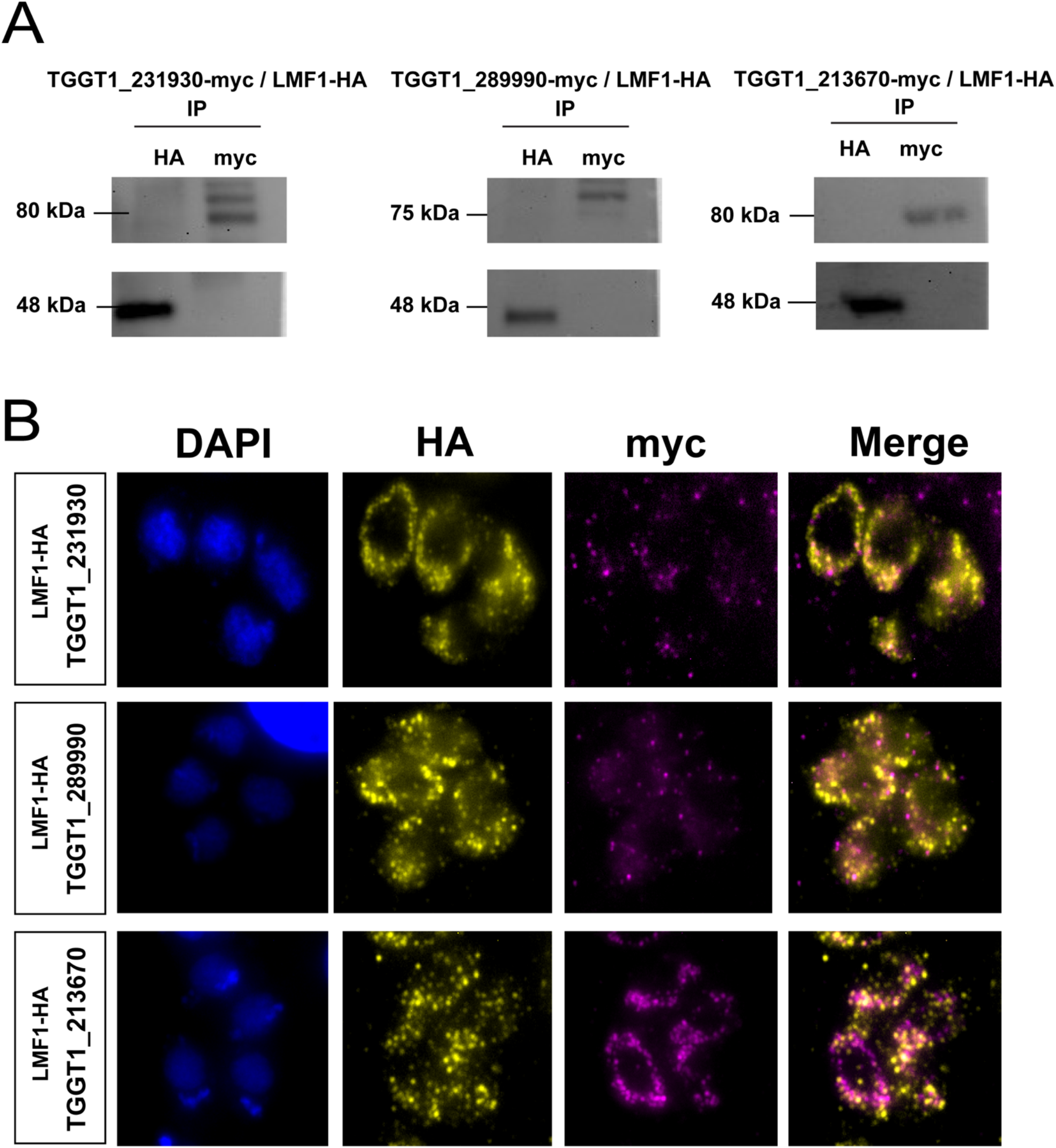
Characterization of LMF1 interactors. To investigate the localization of LMF1 interactors, we introduced sequences encoding an N-terminal Myc tag to the endogenous locus in the parasite strain expressing an HA-tagged LMF1. A) Reciprocal co-immunoprecipitation of putative LMF1 interactors was performed for the strains expressing LMF1-HA and either TGGT1_231930-Myc, TGGT1_289990-Myc, or TGGT1_213670-Myc. For each of the three dually tagged parasite strains, proteins were immunoprecipitated with either anti-HA or anti-Myc conjugated beads and probed with either Myc (for the interactor) and for HA (for LMF1). B) Intracellular parasites expressing the Myc tagged versions of TGGT1_231930-myc, TGGT1_289990-myc and TGGT1_213670-myc were stained for HA (yellow) and myc (magenta).

**Supplemental figure S2.**
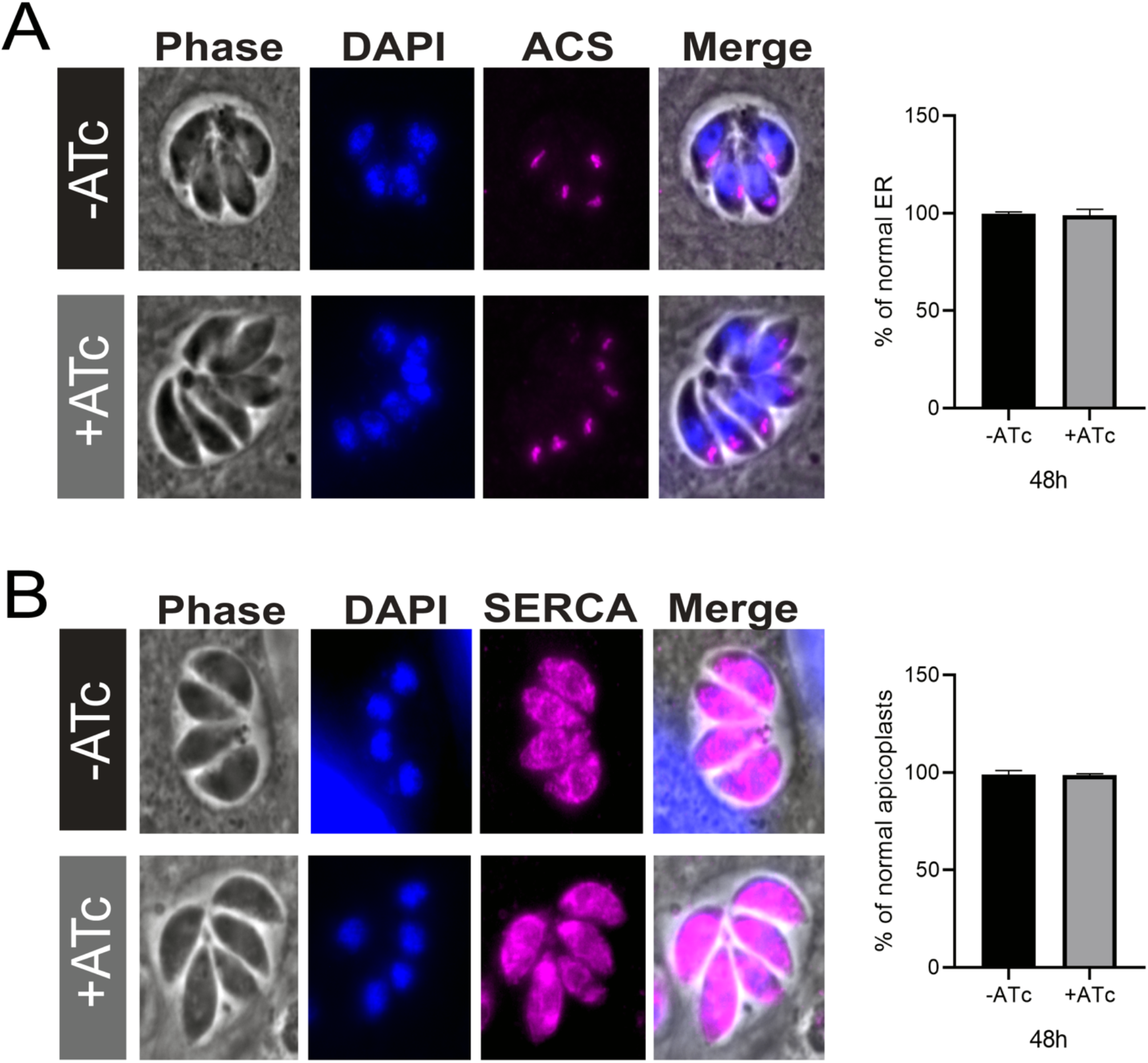
IMC10 knockdown does not affect apicoplast or endoplasmic reticulum (ER) morphology. IFA of parasites stained for DAPI (blue), apicoplast, and ER (magenta) showing the shape of both structures in parasites maintained with and without ATC for 48h. A) Apicoplast morphology at 48h. B) ER morphology at 48h. Scale bar = 5 µm. All graphs represent the percentage of vacuoles with the related phenotype. At least 150 vacuoles per sample were inspected. For all graphs, n = 3. Error bars means SD.

